# CD4^+^ tumor-infiltrating lymphocytes secreting T cell-engagers induce regression of autologous patient-derived non-small cell lung cancer xenografts

**DOI:** 10.1101/2024.05.27.596077

**Authors:** Anaïs Jiménez-Reinoso, Magdalena Molero-Abraham, Cristina Cirauqui, Belén Blanco, Eva M. Garrido-Martin, Daniel Nehme-Álvarez, Carmen Domínguez-Alonso, Ángel Ramírez-Fernández, Laura Díez-Alonso, Ángel Nuñez-Buiza, África González-Murillo, Raquel Tobes, Eduardo Pareja, Manuel Ramírez-Orellana, José L. Rodriguez-Peralto, Irene Ferrer, Jon Zugazagoitia, Luis Paz-Ares, Luis Álvarez-Vallina

## Abstract

Adoptive transfer of tumor-infiltrating lymphocytes (TIL) has shown remarkable results in melanoma, but only modest clinical benefit in other cancers, even after TIL have been genetically modified to improve their tumor homing, cytotoxic potential or overcoming cell exhaustion. The required *ex vivo* TIL expansion process may induce changes in the T cell clonal composition, which could likely compromise the tumor reactivity of TIL preparations and ultimately the success of TIL therapy. A promising approach based on the production of bispecific T cell engagers (TCE) by engineered T cells (STAb-T therapy) improves the efficacy of current T cell redirection strategies against tumor-associated antigens in hematological tumors. We studied the TCRβ repertoire in non-small cell lung cancer (NSCLC) tumors and in *ex vivo* expanded TIL from two unrelated patients. We generated TIL secreting anti-epidermal growth factor receptor (EGFR) x anti-CD3 TCE (TIL^STAb^) and tested their antitumor efficacy *in vitro* and *in vivo* using a NSCLC patient-derived xenograft (PDX) model in which tumor fragments and TIL from the same patient were transplanted into *hIL-2* NOG mice. We confirmed that the standard TIL expansion protocol promotes the loss of tumor-dominant T cell clones and the overgrowth of virus-reactive TCR clonotypes that were marginally detectable in primary tumors. We demonstrated the antitumor activity of TIL^STAb^ both *in vitro* and *in vivo* when administered intratumorally and systemically in an autologous immune-humanized PDX EGFR^+^ NSCLC mouse model, where tumor regression was mediated by TCE-redirected CD4^+^ TIL bearing non-tumor dominant clonotypes.

**SIGNIFICANCE:** Epithelial tumor-derived TIL can be engineered to secrete TCE capable of redirecting T cells bearing non-tumor-dominant clonotypes regardless of their phenotype, which could have broad applications in immunotherapy for solid tumors.

## INTRODUCTION

Adoptive cell therapy (ACT) with tumor-infiltrating lymphocytes (TIL) is based on the isolation and *ex vivo* expansion of intratumoral lymphocytes that are infused back into the patient (1). The diverse and tumor-specific T cell receptor (TCR) repertoire and their ability to localize tumors have classically been considered potential advantages of TIL in cancer treatment (2). TIL-based ACT has shown remarkable results in advanced melanoma (3,4), and although the feasibility of TIL expansion has been demonstrated in other cancer types, only modest clinical benefit has been reported in non-melanoma tumors (5). Current knowledge supports that the *ex vivo* expansion process induces changes in the clonal composition of T cells that are likely to affect the tumor reactivity of TIL preparations (6), as neoantigen reactive T cells (NRTs) may be lost or dramatically under-represented in the expanded TIL products (7).

Genetic modification of TIL has been explored to improve their tumor homing ability, their cytotoxic potential or to overcome T cell exhaustion (8). A promising ACT called STAb-T (<u>s</u>ecreting <u>T</u> cell-engaging <u>a</u>nti<u>b</u>odies), in which T cells are engineered to secrete small bispecific T cell-engagers (TCE), has shown superior efficacy in hematological malignancies (9–11), as the polyclonal recruitment of both engineered and unmodified bystander T cells by secreted TCE leads to a significant boost in antitumor T cell responses.

We hypothesize that the application of STAb-T strategies to TIL products could enhance their therapeutic impact in solid tumors. Intratumoral secretion of TCE by engineered TIL could promote an initial TCR-independent polyclonal response with TCE-mediated T cell activation and cytotoxicity that could lead to a reduction in tumor burden, eventually followed by a second TCR-dependent neoantigen-specific T cell activation and cytotoxicity wave. In this article, we describe for the first time the generation of engineered TIL secreting a bispecific anti-epidermal growth factor receptor (EGFR) x anti-CD3 TCE (TIL^STAb^) and demonstrate its therapeutic potential in an autologous immune-humanized patient-derived xenograft (PDX) of EGFR^+^ non-small cell lung cancer (NSCLC) mouse model.

## MATERIALS AND METHODS

### Tumor and donor PBMC sample collection

Tumor samples from two patients with locally advanced non-small cell lung cancer (NSCLC) were included in this study. Patient 1 was a 76-years old man, diagnosed with squamous cell carcinoma (T4N0M0) in 2018. HLA typing: A*29:01, A*30:02; B*41:01, B*51:01; DRB1*03:01, DRB1*08:04. Patient 2 was a 56-years old man, diagnosed with adenocarcinoma (T4N1M0) in 2017. A portion of freshly resected surgical tumor specimen from each patient was used for the purposes of this study; the remaining tumor sample was formalin-fixed and paraffin-embedded (FFPE) for routine diagnostic purposes. Donor PBMC for allogeneic TIL stimulation during Rapid Expansion Protocol (REP) were obtained from volunteer healthy donors’ buffy coats by density-gradient centrifugation using Lymphoprep^TM^ (Axis-Shield, Oslo, Norway). Alternatively, PBMC for cytotoxicity assays were obtained from volunteer healthy donors’ freshly isolated peripheral blood. All samples were obtained after written informed consent from the donors and all studies were performed according to the principles expressed in the Declaration of Helsinki and approved by the Institutional Research Ethics Committees of the hospitals and research centers involved.

### Tumor sample disaggregation for tumor phenotyping

NSCLC samples were collected in RPMI-1640 medium just after surgery, and were mechanically disaggregated and enzymatically digested with Liberase™ DL and deoxyribonuclease I from bovine pancreas (both from Sigma-Aldrich) for 15 minutes at 37 °C. After digestion, samples were centrifuged and resuspended in PBS pH 7.4 supplemented with 2 mM EDTA and 0.5% (w/v) BSA (Sigma-Aldrich) and filtered through 0.70 µm cell strainers to remove fibers and large aggregates. Finally, alive cells from cell suspension were counted and immunophenotyped (**Supplementary Table S1**) or stored in liquid nitrogen in freezing medium, which consisted in fetal bovine serum (FBS; Sigma-Aldrich, St. Louis, MO, USA) supplemented with 10% (v/v) DMSO (Merck Millipore, Tullagreen, Carrigtwohill, Ireland).

### Tumor cell lines

HEK293T (CRL-3216), CHO-K1 (CCL-61), NALM6 (CRL-3273), Jurkat Clone E6-1 (TIB-152), and MKN45 (CRL-1582, ACC 362) cell lines were purchased either from the American Type Culture Collection (Rockville, MD, USA) or the DSMZ (Braunschweig, Germany). Target cells expressing the firefly luciferase gene (NALM6^Luc^, CHO^Luc^ and MKN45^Luc^) were produced in house. HEK293T and CHO-K1 cells were cultured in Dulbecco’s modified Eagle’s medium (DMEM) (Lonza, Walkersville, MD, USA) supplemented with 2 mM L-glutamine (Life Technologies, Paisley, UK), 10% (v/v) heat-inactivated FBS and antibiotics (100 units/mL penicillin, 100 μg/mL streptomycin) (both from Sigma-Aldrich), referred to as DMEM complete medium (DCM). NALM6, Jurkat and MKN45 cells were cultured in RPMI-1640 (Lonza) supplemented with 2 mM L-glutamine, 10% (v/v) heat-inactivated FBS and antibiotics, referred to as RPMI complete medium (RCM). All the cell lines were grown at 37 °C in 5% CO_2_ and routinely screened for mycoplasma contamination by PCR using the Mycoplasma Gel Detection Kit (Biotools, Madrid, Spain).

### Mice models

Mice were housed at the Animal Facility of the Centro Nacional de Investigaciones Oncológicas (CNIO, Madrid, Spain), where all the *in vivo* experiments were carried out in accordance with the CNIO Ethics Committee of Animal Experimentation. The experimental study protocols were additionally approved by local government (PROEX 084/15 and 253/19). Five to six week-old female nude athymic mice (Athymic Nude-*Foxn1^nu^*) or *hIL-2* NOG mice (NOD.Cg-*Prkdc^scid^ Il2rg^tm1Sug^* Tg(CMV-IL2)4-2Jic/JicTac) were purchased from Envigo (Indianapolis, IN, USA) or Taconic Biosciences (Rensselaer, NY, USA), respectively.

### Vector constructs

The mammalian expression vector pCR3.1-EGa1-(G4S)-OKT3 encoding the anti-EGFR x anti-CD3 LiTE has been previously described (12,13). To generate the lentiviral transfer vector, a synthetic gene encoding the signal peptide OncoM followed by the EGFR V_HH_, a five-residue linker (Gly_4_Ser), the OKT3 scFv (V_H_–V_L_) (14), the human c-Myc epitope tag, and a C-terminal polyHis tag was synthesized by GeneArt AG (ThermoFisher Scientific, Regensburg, Germany). The synthetic DNA was then cloned into the bicistronic vector pCCL-EF1α-CD1a-CAR-T2A-EGFP (15) using *Mlu*I/*BspE*I restriction enzymes, obtaining the plasmid pCCL-EF1α-LiTE-T2A-EGFP.

### Lentivirus production and titration

LiTE-expressing viral particles pseudotyped with vesicular stomatitis virus G (VSV-G) glycoprotein were generated in HEK293T cells using standard polyethylenimine transfection protocols and concentrated by ultracentrifugation, as previously described (9). Viral titers were consistently in the range of 1×10^8^ transducing units/mL. Functional titers of LiTE-encoding lentiviruses were determined by limiting dilution in HEK293T cells and analyzed using green fluorescent protein (EGFP) expression by flow cytometry. Cell acquisition was performed in a FACSCanto™ II flow cytometer using BD FACSDiva software (BD Biosciences, San Jose, CA, USA). Analysis was performed using FlowJo V10 software (Tree Star, Ashland, OR, USA).

### Cell transfection, EGFR LiTE binding and T cell activation assays

HEK293T cells were transfected with the appropriate expression vectors using Lipofectamine 3,000 Reagent (ThermoFisher Scientific, Waltham, MA, USA) according to the manufacturer’s protocol. After 48 hours, transiently transfected HEK293T cells were analyzed by flow cytometry and conditioned media were collected and stored at –20 °C for enzyme-linked immunosorbent assays (ELISA), western blotting and EGFR LiTE binding assays. For EGFR LiTE binding assay, conditioned media from transiently transfected HEK293T cells were incubated with EGFR^‒^CD3^‒^ NALM6, EGFR^‒^CD3^+^ Jurkat or EGFR^+^CD3^‒^ MKN45^Luc^ cells, and analyzed with an APC-conjugated anti-His monoclonal antibody (mAb, Miltenyi Biotec, Bergisch Gladbach, Germany) by flow cytometry. For T cell activation assays, EGFR^‒^ CHO^Luc^ or EGFR^+^ MKN45^Luc^ cells were co-cultured with non-transduced (Jurkat^NTd^) or with EGFR LiTE-transduced (Jurkat^STAb^) Jurkat cells at a 1:1 effector:target (E:T) ratio in the presence of conditioned media from transiently transfected HEK293T cells. After 24 hours, T cell activation was analyzed by flow cytometry using PE-conjugated anti-CD69 mAb. Cell acquisition was performed in a FACSCanto™ II flow cytometer using BD FACSDiva software (BD Biosciences). Analysis was performed using FlowJo V10 software (Tree Star, Ashland, OR, USA).

### T cell transduction

PBMC were plate-coated activated with 1 µg/ml anti-CD3 (OKT3 clone) and 1 µg/ml anti-CD28 (CD28.2 clone) mAbs (BD Biosciences) for 2 days and transduced with EGFR LiTE- encoding lentiviruses (T^STAb^) in the presence of 10 ng/ml interleukin (IL)-7 and 10 ng/ml IL-15 (Miltenyi Biotec). As negative control, non-transduced T cells were used (T^NTd^). T cells were expanded in RCM supplemented with IL-7 and IL-15 (10 ng/mL).

### *In vitro* cytoxicity assays

For cytotoxicity assays, non-transduced (T^NTd^ / TIL^NTd^) or EGFR LiTE-transduced (T^STAb^ / TIL^STAb^) T cells were co-cultured with luciferase-expressing target cells (EGFR^‒^ CHO^Luc^ or EGFR^+^ MKN45^Luc^ cells) at the indicated E:T ratios. After 48 hours or 72 hours, 20 μg/ml D-luciferin (Promega, Madison, WI, USA) was added before bioluminescence quantification using a Victor luminometer (PerkinElmer, Waltham, MA, USA). Percent specific cytotoxicity was calculated using the formula: 100 – [(bioluminescence of each sample*100) / mean bioluminescence of NT-target cells]. Specific lysis was established as 100% of cell viability, and 100% lysis was established by adding 5% (v/v) Triton X-100 into target cells.

### Generation of engineered TIL

TIL were isolated from resected fresh NSCLC samples, washed with Ca^2+^Mg^2+^ free-HBSS (Lonza) and sliced into 1-3 mm^3^ fragments with a sterile scalpel, transferred with the surrounding HBSS into an enzymatic digestion medium containing RPMI-1640, antibiotics, 2mM L-glutamine, 10 µg/mL gentamycin (Lonza), 10 µg/mL dornase alfa (Pulmozyme^®^, Roche, Basel, Switzerland) and 1 mg/mL collagenase IV (Sigma-Aldrich), and incubated for 5 hours at 37 °C. After digestion, cell suspension was filtered through 100 µm cell strainers and washed three times with Ca^2+^Mg^2+^ free-HBSS before counting alive cells and kept in culture (37 °C, 5% CO2) for 14 days in a tissue culture treated 24-well plate (ThermoFisher Scientific) until REP stage in TIL-RCM, which consisted in RPMI-1640 supplemented with 2 mM L-glutamine, antibiotics, 1.25 μg/mL amphotericin B and 25 mM HEPES (both from Lonza), 10% (v/v) heat-inactivated human male AB serum (Sigma-Aldrich) and 6,000 IU/mL recombinant human IL-2 (rhIL-2, Proleukin^®^, Clinigen, Staffordshire, United Kingdom). After a 14-day culture period, TIL were preserved in liquid nitrogen in freezing medium until use. During the pre-REP, TIL were thawed, washed twice and kept in a tissue culture treated 24-well plate until REP stage in TIL-RCM (Resting TIL, Day –5). After two days (Day –3), Pre-REP TIL were activated with plate-coated OKT3 (30 or 300 ng/mL) for 48 hours in tissue culture treated 96-well plates (1×10^5^ cells/well). At day –1, 300 ng/mL OKT3-activated pre-REP TIL were lentivirally transduced in a 20 μg/mL retronectin-coated non-tissue culture treated 96-well plate (16) with LiTE-encoding lentiviruses (TIL^STAb^); 30 ng/mL OKT3-activated pre-REP TIL were kept as non-transduced cells (TIL^NTd^). After 24 hours, pre-REP TIL were expanded for 14 days under a small-scale REP (17): at day 0, pre-REP TIL were co-cultured with 2×10^7^ 40 Gy-irradiated allogeneic feeder PBMC (1:200 ratio) in upright 25-cm^2^ flask (ThermoFisher Scientific) in REP-TIL culture medium, which consisted in a 1:1 mixture of 10 mL of TIL-RCM and 10 mL of CTS™ AIM-V™ SFM medium (ThermoFisher Scientific) supplemented with 1.25 μg/mL amphotericin B, 10% (v/v) heat-inactivated human male AB serum and 6,000 IU/mL rhIL-2. On day 5, 10 mL of culture supernatant was replaced with 10 mL of fresh REP-TIL medium; on day 7 and every two day thereafter, REP TIL were split into further 25-cm^2^ flasks and maintained at 1×10^6^ cells/mL with fresh REP-TIL medium until final REP stage (14 days).

### *In vivo* autologous treatment of *hIL-2* NOG PDX mice

Tumor fragments from P1 (18) were implanted into nude female athymic mice (Athymic Nude-*Foxn1^nu^*, Envigo, Indianapolis, IN, USA) as previously described (Yagüe P. el al, in press). When tumor size reached 800-1000 mm^3^, the animals were euthanized and 5x5 mm tumor fragments were subcutaneously (s.c). implanted into female *hIL-2* NOG [NOD.Cg-*Prkdc^scid^ Il2rg^tm1Sug^* Tg(CMV-IL2)4-2Jic/JicTac, Taconic Biosciences, Rensselaer, NY, USA]. When tumor size reached 100-300 mm^3^ approximately, mice were randomized and autologous TIL^NTd^ or TIL^STAb^ were intratumorally (i.t., 3×10^6^ cells/ mouse); or intravenously (i.v., 7.5×10^6^ cells/ mouse) injected in PBS supplemented with 300 IU/mL rhIL-2 (19). Mice were monitored daily and weighted once per week to detect possible signs of graft versus host disease (GvHD), and tumor size was measured twice a week with a caliper using the following formula: V = (L×W^2^)/2, where L represents the length or largest longitudinal tumor diameter, and W is the width or largest transverse tumor diameter. When tumors grew over 1,000 mm^3^ or GvHD signals appeared, mice were euthanized and tumors, spleens, livers and blood samples were collected.

### Immunohistochemistry and *Alu-II in situ* hybridization

Primary tumors and tumors from different treatment groups were collected and fixed in neutral buffered formalin (Sigma-Aldrich) for 48 hours and after extensive washing in PBS, tissues were embedded in paraffin (FFPE). Four-µm-thick FFPE sections were incubated with the antibodies listed in **Supplementary Table S1** on a Bond^TM^ Automated System (Leica Microsystems, Wetzlar, Germany), according to the manufacturer’s instructions. Alternatively, Discovery ULTRA (Ventana, Roche) or Autostainer Link 48 (Dako, Agilent Technologies, Santa Clara, CA, USA) automated immunostaining platforms were used.

Nuclei were counter stained with Harris’ hematoxylin. Discovery ULTRA (Ventana, Roche) automatic platform was used to run *in situ* hybridization for detecting *Alu* repeated sequences. After deparaffinization, re-hydratation and antigen retrieval, slides were incubated with the DNP-labeled Alu positive control probe II (Ventana, Roche). After washing, slides were incubated with a rabbit anti-DNP and HRP-conjugated OmniMap anti-rabbit antibody (Ventana, Roche) was used for visualization. DAB substrate was used as a chromogen (ChromoMap DAB, Ventana, Roche) and nuclei were counterstained using Carazzi’s hematoxylin; all the slides were dehydrated, cleared and mounted with a permanent mounting medium for microscopic evaluation. Whole digital slides were acquired with a slide scanner (AxioScan Z1, Zeiss, Oberkochen, Germany), and positive versus total cells were automatically quantified using Zen 3.1 Blue or AxioVision 4.6 softwares (Zeiss).

### Digital Spatial Profiling

Protein-based digital spatial profiling (DSP) assay was performed as previously described (20,21). Briefly, FFPE tissue slides derived from the intact primary tumor (one section) and PDX samples treated with TIL^STAb^ (TIL^STAb^: two serial sections) or TIL^NTd^ (TIL^NTd^: two serial sections) were subjected to antigen retrieval procedures with citrate buffer pH 6 (Sigma Aldrich) at 100-112 °C for 15 minutes in a TintoRetriever Pressure Cooker (BioSB, Santa Barbara, CA, USA), followed by 1 hour incubation with Buffer W blocking solution (NanoString Technologies, Seattle, WA, USA). Subsequently, slides were co-incubated overnight with fluorescent-labeled antibodies (morphology markers) to detect tumor cells (pan-cytokeratin or PanCk) and all immune cells (CD45), together with 42-plex photocleavable oligonucleotide-labeled primary antibodies panels (profiling antibodies, **Supplementary Table S2**). In addition, serial tissue sections derived from PDX samples treated with TIL^STAb^ cells and TIL^NTd^ cells (one section each) were stained with fluorescent-labeled antibodies (**Supplementary Table S1**) to detect tumor cells (PanCk), CD4 T cells (Abcam, Cambridge, UK), and CD8 T cells (Novus Biologicals, Centennial, CO, USA) together with the same 42-plex photocleavable oligonucleotide-labeled primary antibody panels. The next day, a post-fixation step using 4 % paraformaldehyde (ThermoFisher Scientific) for 30 minutes onto the slides was followed by nuclear staining with SYTO13 (NanoString Technologies) for 15 minutes. Once the staining was completed, the slides were loaded on the GeoMx DSP instrument (NanoString Technologies), where they were scanned to produce a digital fluorescent image of the tissue. Next, we generated multiple circular regions of interest (ROI) of a maximum of 500 µm from PanCk^+^ intratumoral areas in each slide (**Fig. 1D**). In addition, to obtain 42-plex protein measurements selectively and separately from CD4 and CD8 compartments, we generated molecularly defined CD4^+^ and CD8^+^ compartments by fluorescent marker colocalization in each of the ROI from the PDX samples treated with TIL^STAb^ cells and TIL^NTd^ cells (**Fig. 3C**). Subsequently, the oligos from these ROI of CD4/CD8 specific compartments were released upon UV-light exposure, collected via microcapillary aspiration and dispensed into a 96-well plate. Then, these oligos were hybridized to 4-color, 6-spot optical barcodes and digitally counted in the nCounter system (NanoString Technologies). Using the GeoMx software (NanoString Technologies), digital counts were first normalized to internal spike-in controls (ERCCs) to account for system hybridization variation and then normalized to the counts of two housekeeper protein probes (GADPH and S6).

**Figure 1.**
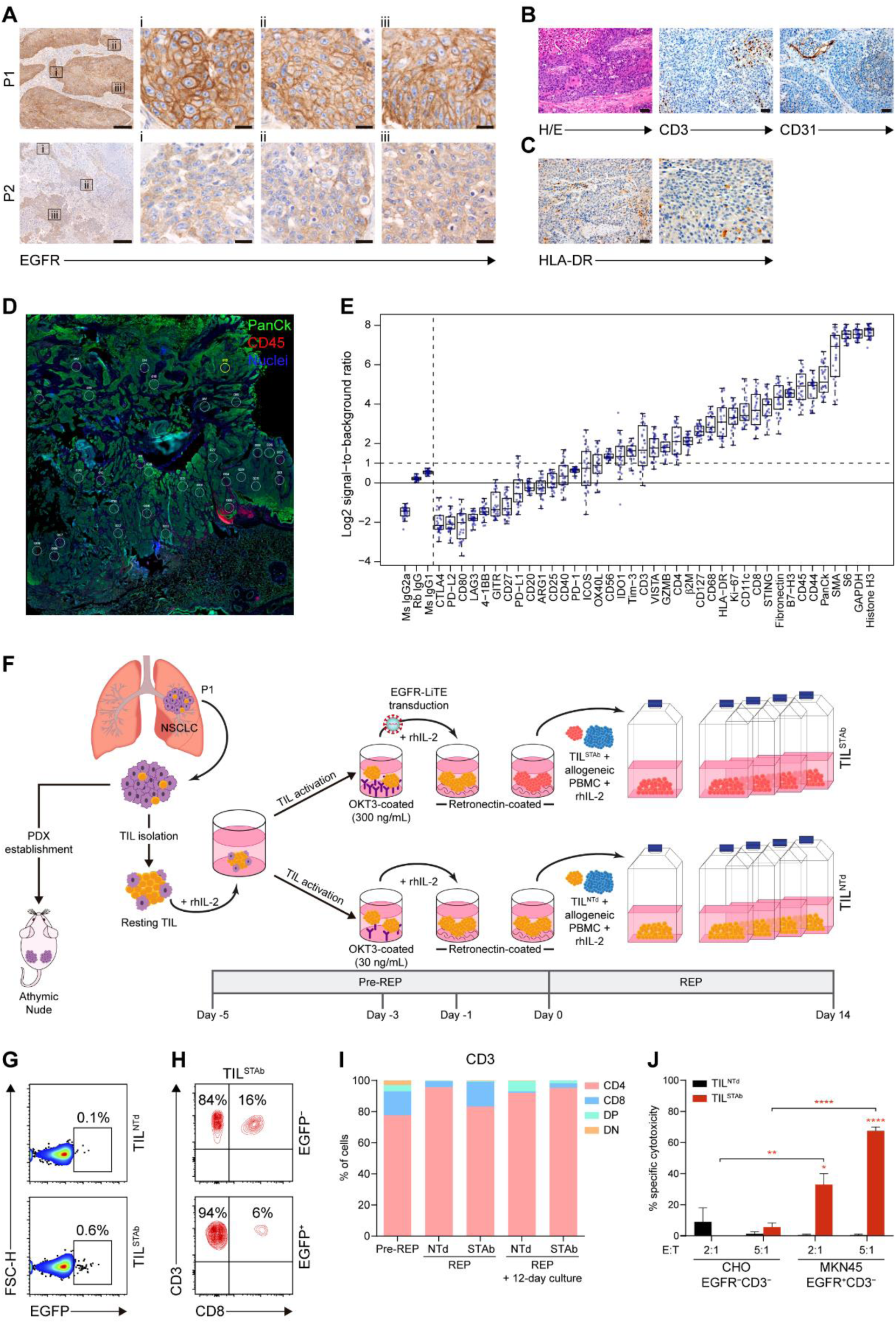
Phenotype of primary tumors and generation of patient-derived engineered EGFR-TCE-secreting TIL. **(A)** EGFR expression by IHC in P1 and P2 primary tumors. Magnification of three ROI (i, ii, iii) from left images are shown. Scale bars: 200 µm and 20 µm in magnified images. **(B)** Histological detail of the P1 primary tumor (H/E), T cell infiltration (CD3^+^) and vascularization (CD31^+^) by IHC. The P1 tumor was a pleomorphic squamous cell carcinoma with atypical dyskeratotic cells and large necrotic areas. Scale bars: 50 µm. **(C)** HLA-DR expression by IHC in P1 primary tumor. Scale bars: 50 µm (left) and 20 µm (right). **(D)** DSP image showing the multiple circular ROI of a maximum of 500 µm from PanCk^+^CD45^+^ intertumoral areas in P1 primary tumor. **(E)** Signal-to-noise plot showing the log2-transformed signal (counts of each protein) relative to background levels (averaged counts of the three negative controls) of all markers included in the DSP immune-related protein panel in P1 primary tumor sample. **(F)** Engineered EGFR-STAb TIL (TIL^STAb^) small-scale REP generation and PDX establishment from P1 NSCLC primary tumor. **(G)** Percentage of reporter EGFP expression in non-transduced TIL (TIL^NTd^) and TIL^STAb^ after REP. **(H)** Percentages of CD4^+^ and CD8^+^ T cells within EGFP^-^ and EGFP^+^ TIL^STAb^ cells. **(I)** Percentages of CD4^+^, CD8^+^, double positive (DP) and double negative (DN) T cells among pre-REP, REP and 12 days after REP cultured TIL^NTd^ and TIL^STAb^ cells. **(J)** Specific cytotoxicity 12 days after REP of TIL^NTd^ or TIL^STAb^ cells against EGFR-negative (CHO) or EGFR-positive (MKN45) cells at the indicated E:T ratios after 72 hours. The percentage of specific cytotoxicity was calculated by adding D-luciferin to detect bioluminescence. Data represent mean+SEM and statistical significance was calculated by two-way ANOVA test corrected by Tukey’s multiple comparisons test; *p < 0.05; **p < 0.01, ****p < 0.0001. TCE, bispecific T cell-engagers; IHC, Immunohistochemistry; P1, Patient 1; P2, Patient 2; ROI, region of interest; H/E: hematoxylin and eosin; REP, rapid expansion protocol; DSP, digital spatial profiling; SEM, standard error of mean.

### Enzyme-linked immunosorbent assays

To detect the EGFR LiTE secreted to culture supernatants, 2.5 µg/mL recombinant human EGFR Fc chimera protein (R&D Systems, Minneapolis, MN, USA) was immobilized on Maxisorp plates (ThermoFisher Scientific) overnight at 4 °C. After washing and blocking with PBS supplemented with 5% (w/v) BSA (Sigma-Aldrich), conditioned media were added and incubated at room temperature for 1 hour. Then, wells were washed 3 times with PBS supplemented with 0.05% (v/v) Tween^®^20 (Sigma-Aldrich) and 3 times with PBS, and anti-His mAb (QIAGEN, Hilden, Germany) was added (1 µg/mL in PBS supplemented with 1% w/v BSA). After washing, horseradish peroxidase (HRP) AffiniPure goat anti-mouse IgG (H+L) (0.4 µg/mL in PBS supplemented with 1% w/v BSA; Jackson ImmunoResearch, West Grove, PA, USA) was added, and the plate was developed using tetramethylbenzidine (Sigma-Aldrich). Human IL-2 secretion in *hIL-2* NOG PDX mice treated with TIL^NTd^ or TIL^STAb^ cells was analyzed by ELISA (Diaclone, Besancon Cedex, France), following manufacturer’s instructions.

### Western blotting

Samples were separated under reducing conditions on 10-20% Tris-glycine gels (Life Technologies), transferred onto PVDF membranes (Merck Millipore) and probed with 0.5 µg/mL anti-Myc mAb, clone 9E10 (Merck Millipore), followed by incubation with 0.8 µg/mL HRP-conjugated goat anti-mouse IgG (Fc specific, Sigma-Aldrich). Visualization of protein bands was performed with Pierce™ ECL Plus Western Blotting Substrate (ThermoFisher Scientific) and ChemiDoc™ MP Imaging System machine (Bio-Rad Laboratories, Hercules, CA, USA).

### TCRβ CDR3 repertoire sequencing

DNA from TIL samples was isolated using QIAamp DNA Blood Mini Kit (QIAGEN). DNA from FFPE samples was isolated using RNA/DNA Purification Plus Kit (Norgen Biotek, Thorold, Ontario, Canada). To obtain the sequence of the most abundant TCRβ CDR3 clonotypes, next-generation sequencing (NGS) was applied from either FFPE TIL-treated tumor samples or from pre/post-REP TIL DNA samples. DNA was amplified in 3 PCR steps (PCR1, PCR2 and PCR3), using the SuperPlex Premix PCR master mix (Takara, Otsu, Shiga, Japan). PCR1: multiplex PCR (touchdown approach) using 39 Forward V sequences primers and 13 Reverse J sequences primers. PCR2: to incorporate sample barcodes and Illumina sequencing primer and library adapters (p5 and p7) with the products of the first PCR. PCR3: was done with p5 and p7 primers to ensure that the entire amplicon product was able to be sequenced. After each PCR step, agarose gel electrophoresis were performed in order to isolate amplified sequences of the expected size and to prevent having primers or primer dimers (PCR1 and PCR2) or for isolating each sample’s DNA library for sequencing. Finally, prepared libraries were sequenced on the Illumina MiSeq V2 300 cycles (single reads) platform.

### TCRβ CDR3 repertoire analysis

TCRβ V(D)J gene usage, CDR3 sequences identification and clonotype information were determined using IgBLAST (https://www.ncbi.nlm.nih.gov/igblast/), which defines a clonotype as any V(D)J rearrangements with the same germline V(D)J gene segments, the same productive/non-productive status and the same CDR3 nucleotide as well as amino sequence. Those having the same CDR3 nucleotide but different amino acid sequence or productive/non-productive status due to frameshift in V or J gene are assigned to a different clonotype. All productive clonotypes with same amino acid sequence but different nucleotide sequence were clustered based on their amino sequence. Given that the cysteine (C) at the beginning and the phenylalanine (F) at the end of the CDR3 are conserved in TCRβ,

IgBLAST does not include these positions in their results; thus, all clonotypes shown in this paper exclude the C and the F 5’ and 3’ positions, respectively. TCRβ CDR3 repertoire metrics and statistics are detailed in **Supplementary Table S3**. For TCR diversity calculation, we used the Shannon’s [1] and the Gini-Simpson’s [2] diversity indexes. TCR clonality (C) was calculated as 1 – Pielou’s evenness [3]. In [1] and [2], Σ = sum from clone 1 to clone N; N = total number of productive unique clonotypes with 5-20 amino-acid length; *pi* = proportion of the sequence of our interest in the entire population.

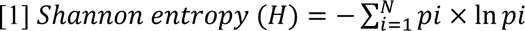

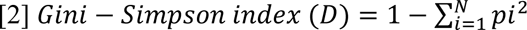

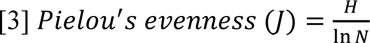

Plots were generated using GraphPad Prism 6 (GraphPad Software Inc., La Jolla, CA, USA). Venn diagrams were generated using InteractiVenn (22).

### Flow cytometry

Antibodies and other reactives used for immune phenotyping of primary tumor samples and *ex vivo* TIL cultures analyzed by flow cytometry are detailed in **Supplementary Tables S1** and **S2**. Disaggregated tumor samples were thawed at 37 °C and immediately resuspended in PBS (Sigma-Aldrich) to remove traces of DMSO. All staining procedures of disaggregated tumors started with Fc receptor Blocking Reagent incubation (1 µl/ 1×10^6^cells) (Miltenyi Biotec) for 10 minutes at 4 °C, followed by cell viability staining with LIVE/DEAD™ Fixable Blue/Violet (2 µl/ 1×10^6^cells) (Invitrogen, ThermoFisher Scientific) for 20 minutes at room temperature in darkness. Different antibodies were used to check tumor infiltrating T, B and NK cells and macrophages (**Supplementary Table S1**). After incubation, samples were washed and resuspended in PBS. Cell surface-bound LiTE TCE were detected using an APC-conjugated anti-His mAb (Miltenyi Biotec). Cell acquisition was performed using the BD FACSCanto™ II or BD LSRFortessa™ flow cytometers using BD FACSDiva software (BD Biosciences). Analysis was performed using FlowJo V10 software.

### Quantification and statistical analysis

Statistical tests were performed using GraphPad Prism 6/8 (GraphPad Software Inc.) software. Results of experiments are expressed as mean, mean ± standard deviation (SD) or mean ± standard error of the mean (SEM). Significance was considered only when p values were less than 0.05 (*p < 0.05; **p < 0.01; ***p < 0.001, ****p < 0.0001). For DSP analysis (GeoMx software), non-parametric tests (Benjamini-Hochberg false discovery rate method) comparing ROI-based or compartment-based DSP protein counts between tumor samples were used. To analyze differentially expressed proteins, the fold change of protein levels in ROI/compartments from TIL^STAb^-treated relative to TIL^NTd^-treated tumor samples was quantified. To assess immune microenvironment features in all intratumoral regions, each ROI or compartment was considered as an independent sample. All hypothesis testing was performed at a two-sided significance level of α=0.05.

### Data availability

All data relevant to the study are included in the article or uploaded as supplementary information. Further information and requests for resources and reagents should be directed to and will be fulfilled by the lead contact, Luis Álvarez-Vallina. Plasmids generated in this study will be made available on request, but we may require a completed Material Transfer Agreement if there is potential for commercial application. This paper does not report original code. Any additional information required to reanalyze the data reported in this paper is available from the lead contact upon request.

## RESULTS

### *Ex vivo* culture modifies T cell phenotype and TCR repertoire

Primary NSCLC tumor samples from two patients (P1 and P2) were processed, and TIL were isolated and expanded under a small-scale rapid expansion protocol (REP) (17). Analysis of the TCRβ clonotype composition in primary tumors and in *ex vivo* TIL cultures (**Supplementary Table S1**) revealed some differences between samples from P1 and P2. While P1 primary tumor (**Supplementary Fig. S1A**) shared more clonotypes with *ex vivo* cultured TIL than P2 (**Supplementary Fig. S1B**), further analysis of the 30-most frequent (TOP-30) clonotypes confirmed that *ex vivo* TIL expansion markedly reduced the TCRβ repertoire diversity found in primary tumors, but increased the repertoire clonality (6) (**Supplementary Fig. S1C-E**), with one predominant clonotype (ASSDGRYGYT) in P1 and two overrepresented clonotypes (ASSRDGQGQGEQF and ASSSGLYGYT) in P2 (**Supplementary Fig. S1F** and **G**).

### Phenotypic characterization of NSCLC tumors

Disaggregated P1 and P2 samples differed in their CD4^+^ and CD8^+^ TIL composition, as P1 TIL became predominantly CD4^+^ cells after *ex vivo* expansion (**Supplementary Fig. S2A** and **B**). Immunohistochemical (IHC) analysis of primary tumors revealed a strong and homogeneous cell membrane EGFR staining in P1 (**Fig. 1A**, top panels) and a very weak EGFR staining in P2 (**Fig. 1A**, bottom panels), but no *EGFR* mutations were detected in any tumor.

Since we aim to study a TIL-redirecting strategy targeting EGFR-expressing tumors, we focused on P1 tumor. Mutational analysis revealed missense (D1017H, M1043I) and synonymous (X1069X) changes in *PI3KCA*, a missense mutation in *CDKN2A* (A20P) and a splicing mutation in *TP53* (c.523+1G>T), with a TMB of 9 (23). Isolated CD31^+^ vascular structures and sparse infiltration of CD3^+^ cells were observed between neoplastic cell nets (**Fig. 1B**). HLA-DR IHC analysis revealed the presence of inflammatory infiltrate but not HLA-DR^+^ tumor cells (**Fig. 1C**). Digital Spatial Profiling (DSP) of immune-related targets (**Fig. 1D**, showing different regions of interest or ROI) confirmed that P1 tumor was infiltrated by CD45^+^ cells, with similar proportions of CD8 and CD4, and low or negative CTLA-4, PD-L1, LAG-3 and 4-1BB expression (**Fig. 1E**), as shown after tumor disaggregation (**Fig. S2C**).

### Engineered STAb-T cells efficiently secrete functional EGFR-specific TCE and induce specific cytotoxic responses

The small (≈43 kDa) anti-EGFR x anti-CD3 <u>li</u>ght <u>T</u> cell <u>e</u>ngager (EGFR LiTE) (12,13) was cloned under the control of the *EF1α* promoter into a T2A-based bicistronic lentiviral vector (**Supplementary Fig. S3A** and **B**) and was secreted in functional active form (**Supplementary Fig. S3C-E**). Jurkat T cells were efficiently transduced (Jurkat^STAb^) according to the percentages of EGFP^+^ cells and surface-bound EGFR LiTE (cell decoration) (**Supplementary Fig. S3F** and **G**) and were specifically activated after 24-hour co-culture with EGFR^+^ cells, but no T cell activation was detected in co-cultures with EGFR^‒^ or with non-transduced Jurkat cells (Jurkat^NTd^) (**Supplementary Fig. S3H** and **I**). To study the cytotoxicity potential, primary T cells transduced with EGFR LiTE-encoding lentivirus (T^STAb^) or non-transduced (T^NTd^) (**Supplementary Fig. S3J**), were co-cultured with EGFR^‒^ and EGFR^+^ cells at different effector:target (E:T) ratios. T^STAb^, but not T^NTd^ cells, specifically killed EGFR^+^ cells, reaching 70% and 90% cytotoxicity at 2:1 and 5:1 E:T ratios, respectively (**Supplementary Fig. S3K**).

### Generation of engineered STAb-TIL

To investigate the therapeutic potential of P1-derived standard TIL and engineered EGFR LiTE secreting TIL, a modified small scale-REP (16) was performed with lentiviral transduction prior to the allogeneic PBMC co-culture (**Fig. 1F**). Cell growth was monitored on days 7, 9, 12 and 14, and no differences were observed between non-transduced and EGFR LiTE-transduced TIL, TIL^NTd^ and TIL^STAb^ respectively (**Supplementary Fig. S4A**). Transduction efficiency after REP was **≈**1% (**Fig. 1G**), and pre-REP and REP phenotype analysis demonstrated that *ex vivo* expanded TIL were biased towards the effector memory (EM) CD4^+^ T cell compartment, whereas the CD8 compartment expressed higher levels of exhaustion markers and was predominantly composed of central memory (CM) cells (**Fig. 1H** and **I** and **Supplementary Fig. S4B-E**). Twelve days after REP, TIL^NTd^ and TIL^STAb^ CD4 bias increased (**Fig. 1I**), but only TIL^STAb^ were able to specifically kill EGFR^+^ cells *in vitro* after 72-hour co-culture with tumor cells (**Fig. 1J**).

### *In vivo* treatment with autologous engineered TCE-secreting TIL

To study the antitumor activity of TIL^STAb^, an autologous NSCLC PDX model was established. Previous findings reported that continuous presence of human interleukin-2 (IL-2) is necessary for tumor eradication by melanoma TIL in NOG mice (19). Therefore, P1 tumor fragments were subcutaneously (s.c.) implanted into *hIL*-2 NOG transgenic mice and TIL^NTd^, TIL^STAb^ or vehicle were injected intratumorally (i.t.) (**Supplementary Fig. S4F**) or intravenously (i.v.) (**Fig. 2A**). TIL^STAb^ injected i.t. were more effective in controlling tumor growth (**Fig. S4G** and **H**), where no human IL-2 consumption was observed (**Supplementary Fig. S4I**). Tumor growth control was similar between i.v. TIL^NTd^ and TIL^STAb^ (**Fig. 2B** and **Supplementary Fig. S4J**), but a significant reduction in human IL-2 plasma levels was observed in TIL^STAb^-treated mice (**Fig. 2C**), suggesting IL-2 consumption in this context. IHC tumor staining from i.v.-TIL^STAb^-treated mice showed increased infiltration of CD3^+^ T cells (**Fig. 2D** and **E**), and one mouse exhibited complete eradication of the NSCLC EGFR^+^ cells (**Fig. 2D**, M17).

**Figure 2.**
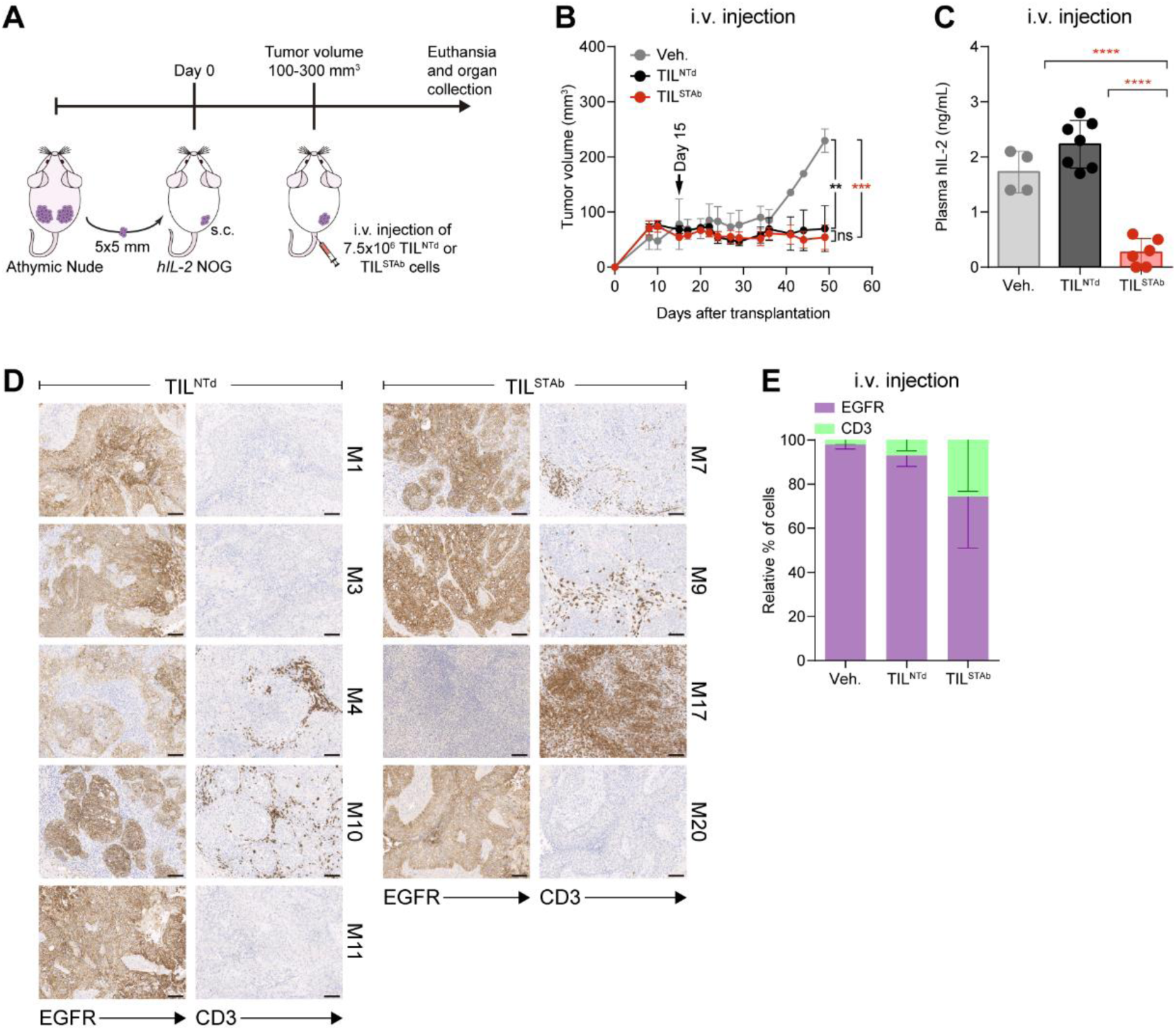
Treatment of EGFR^+^ NSCLC PDX mouse model with autologous engineered TCE-secreting TIL. **(A)** Experimental design. Fifteen *hIL-2* NOG mice were xenografted with P1-derived PDX fragments and when the average tumor volume reached approximately 100-300 mm^3^, were randomized into three groups (2/7/6) and treated i.v. with Veh., (PBS supplemented with 300 IU/mL rhIL-2, *n*=2 mice), TIL^NTd^ (*n*=7 mice) or TIL^STAb^ (*n*=6 mice) (7.5×10^6^ cells/mouse). **(B)** Tumor growth curves shown as mean±SEM; black arrow indicates the day of the i.v. injection. **(C)** Human IL-2 plasma levels in mice after treatment (mean±SD). **(D, E)** EGFR expression and T cell infiltration after TIL treatment by IHC. Scale bars: 100 µm. Data in **(E)** represent mean‒SEM; no significant differences were found. Significance was calculated by two-way **(B,E)** or one-way **(C)** ANOVA test corrected by Tukey’s multiple comparisons test. ns, non-significant; **p < 0.01, ***p < 0.001; ****p < 0.0001. PDX, patient-derived xenograft; Veh., vehicle; IHC, immunohistochemistry; SD, standard deviation; SEM, standard error of mean; M, mouse.

### Tumor regression is mediated by CD4 T cells

To further investigate the mechanisms of tumor regression after systemic TIL^STAb^ therapy in M17, additional markers were analyzed by IHC and DSP and compared with those found in the most CD3^+^ T cell-infiltrated tumor sample from the TIL^NTd^-treated mice (**Fig. 2D**, M4), revealing an increased human CD4^+^ T cell infiltration in M17 mouse with substantial cytotoxic activity (**Fig. 3A** and **B**), which was not associated with detectable HLA-DR expression (**Supplementary Fig. S5A**). While TCE-secreting TIL represented 1% of the total TIL^STAb^ cells before injection (**Fig. 1G**), an increased number of intratumoral EGFP^+^ T cells was detected in M17 mouse (**Fig. 3A** and **B**). DSP analysis confirmed that TIL^STAb^-treated M17 mouse tumor had a significantly higher CD4^+^ T cell infiltration, with immunomodulatory and cytotoxicity markers upregulated in both non- and CD4/CD8- segmented analysis (**Fig. 3C-E** and **S5B** and **C**). Furthermore, co-stimulatory (4-1BB and ICOS) and cytotoxicity (granzyme B) markers correlated better with infiltrating CD4 T cells in TIL^STAb^-treated M17 mouse tumor, but not in P1 primary tumor or in TIL^NTd^-treated M4 mouse tumor, where such correlation was better associated with infiltrating CD8 T cells (**Figure 3F**), suggesting that CD4^+^ TIL were driving the cytotoxic activity (24).

**Figure 3.**
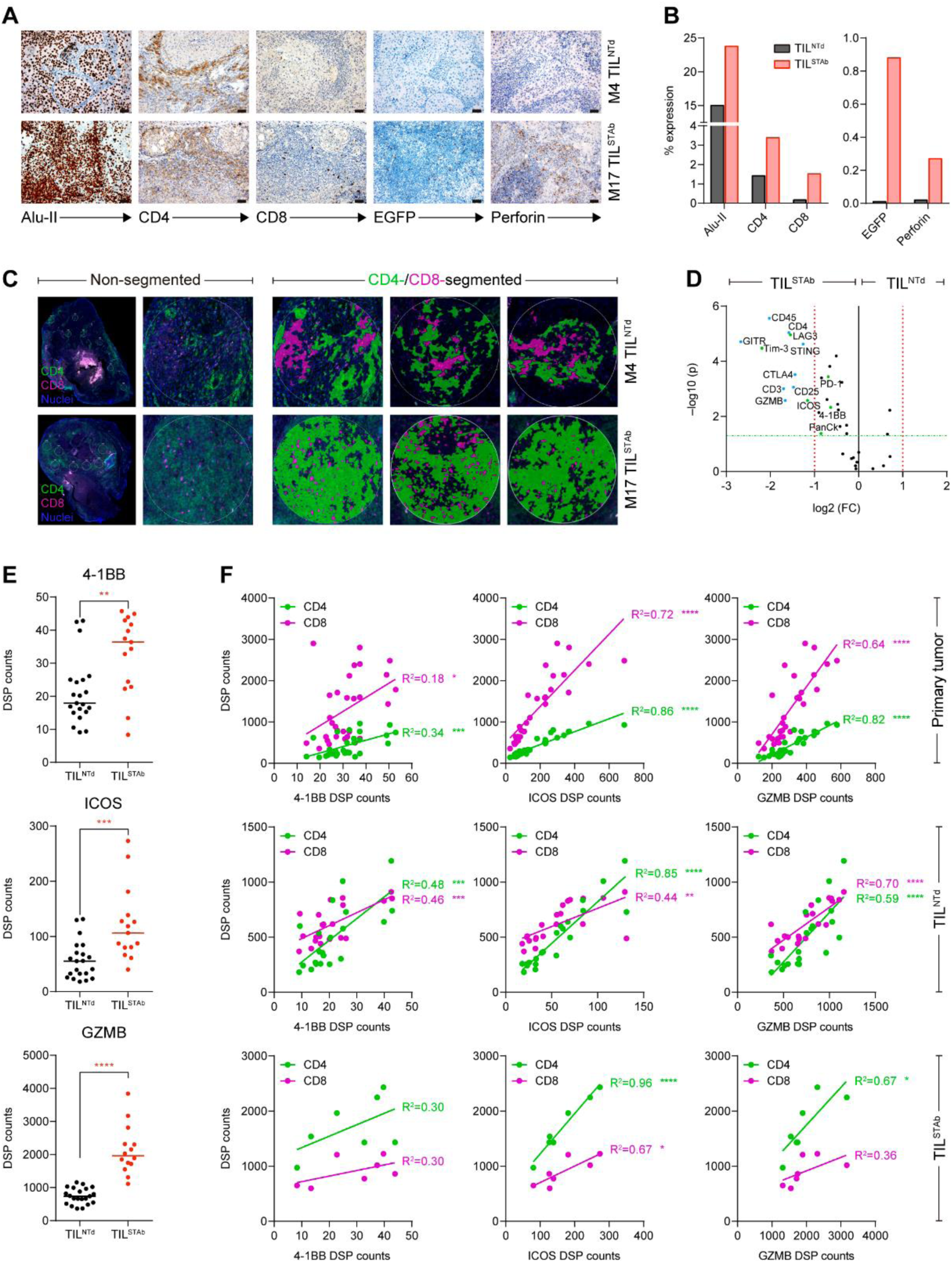
Immunohistochemistry and DSP analysis of intravenously treated NSCLC xenografted *hIL2* NOG mouse 4 (TIL^NTd^-treated) and mouse 17 (TIL^STAb^-treated). (A, B) Expression **(A)** and quantification **(B)** of Alu-II, CD4, CD8, EGR and perforin after treatment of *hIL-2* NOG xenografted mouse 4 (M4) with TIL^NTd^ or mouse 17 (M17) with TIL^STAb^. Scale bars: 50 µm. **(C)** CD4 and CD8 T cell distribution by DSP from multiple circular ROI of a maximum of 500 µm within TIL^NTd^- or TIL^STAb^-treated tumors in non-segmented or CD4/CD8-segmented compartments. **(D)** Differentially expressed protein markers in ROI from tumors treated with TIL^NTd^ or TIL^STAb^ within the lymphocyte (PanCk^‒^ CD45^+^) compartment. Significance (FDR-adjusted p-values) is expressed relative to the FC in protein levels in TIL^STAb^-treated vs. TIL^NTd^-treated ROI. Markers with a FC > 2 and FDR- adjusted p values < 0.05 in the two cohorts (TIL^STAb^- or TIL^NTd^-treated) are marked in bold. **(E)** Comparative analysis of 4-1BB, ICOS and GZMB levels measured by DSP in non- CD4/CD8-segmented or CD4 and CD8 compartments. Significance in **(E)** was calculated by unpaired t-test. **(F)** Linear regression analysis between CD4 or CD8 infiltrated T cells and 4- 1BB, ICOS and GZMB in P1 primary tumor, TIL^NTd^-treated M4 and TIL^STAb^-treated M17 measured by DSP. R^2^ represents goodness of fit to the model. *p < 0.05; **p < 0.01; ***p < 0.001, ****p < 0.0001. FC, fold change; FDR, false discovery rate; ROI, regions of interest; GZMB, granzyme B.

### Tumor regression is mediated by *ex vivo* clonally expanded TIL

Comparison of TCRβ clonotypes from i.v. TIL^NTd^- or TIL^STAb^-treated mouse tumors with the primary tumor and *ex vivo* products (**Fig. 4** and **Supplementary Fig. S6**) showed a restoration of clonotype diversity and a decrease in clonality (**Supplementary Fig. S6A**), with clonotypes similar in CDR3 length to those observed in the primary tumor (**Fig. 4A** and **B**). Within the TOP-30 clonotypes, the *ex vivo* expanded ASSDGRYGYT clonotype remained the most prominent in the TIL^STAb^-treated M17 mouse tumor (**Fig. 4C**, red-colored slices), indicating that tumor regression was driven by the redirection of CD4^+^ REP TIL by locally secreted EGFR-TCE. Furthermore, only TIL^STAb^-treated tumors shared a common clonotype with the primary tumor, which was the *ex vivo* expanded ASSDGRYGYT (**Supplementary Fig. S1F, S6B** and **C**).

**Figure 4.**
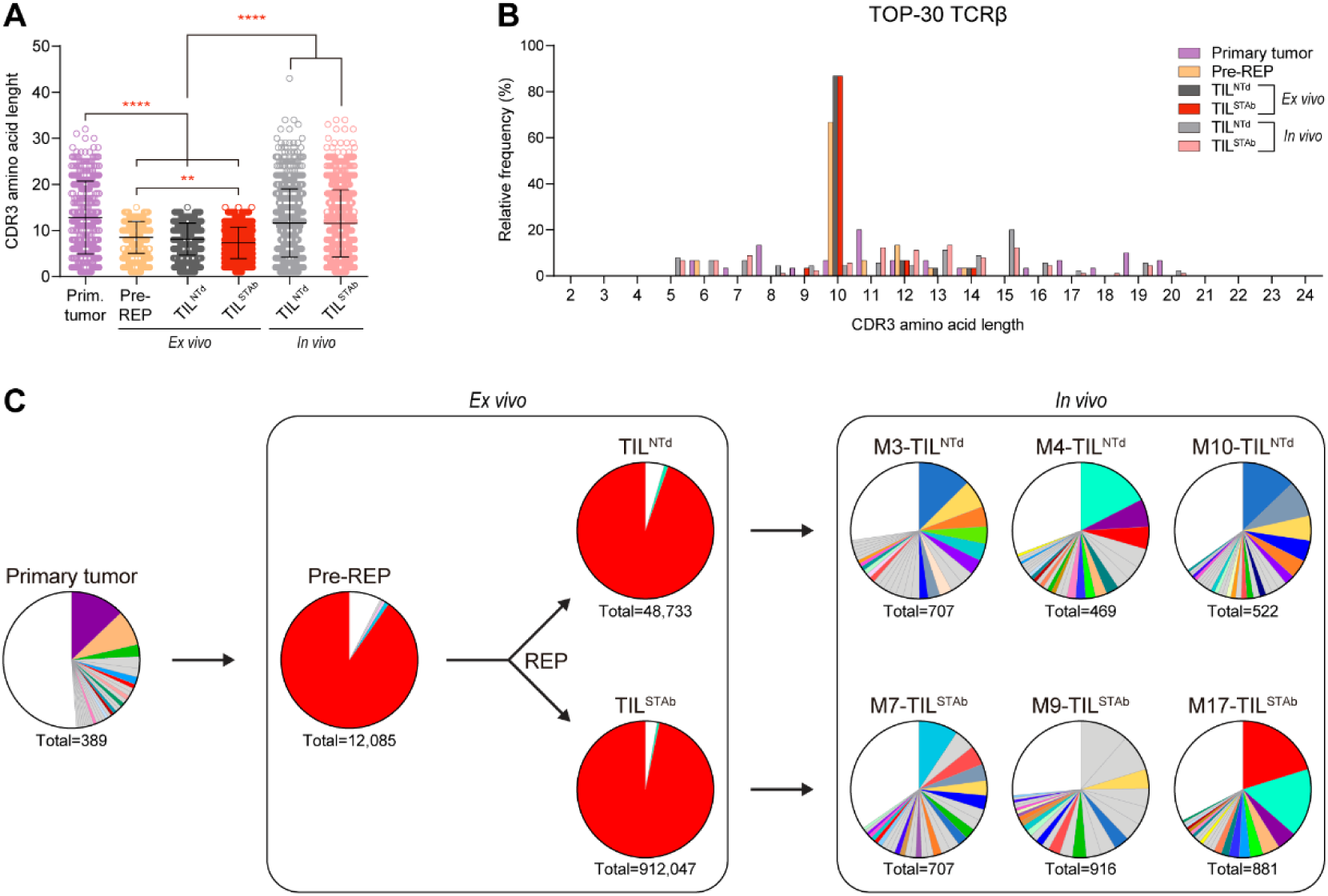
TCR clonotypes composition of primary tumor, *ex vivo* TIL and *in vivo* TIL- treated tumors from P1. **(A)** TCRβ CDR3 length comparison of total unique productive clonotypes. Data represent mean±SD and statistical significance was calculated by one-way ANOVA test corrected with a Tukey’s multiple comparisons test; **p < 0.01; ****p < 0.0001. **(B)** TCRβ CDR3 length distribution of the TOP-30 productive clonotypes. **(C)** Frequency of TOP-30 productive clonotypes. Only clonotypes with frequencies above 0.5% are represented. Shared colors among different pie charts represent clonotypes with identical amino acid sequence. Numbers below pie charts represent total reads assigned to unique productive clonotypes. Only clonotypes with 5-20 amino acid length CDR3 are included in **(B-C)**.

## DISCUSSION

Here, we describe the first ACT strategy using TIL secreting TCE (TIL^STAb^) and demonstrate their enhanced therapeutic activity over standard TIL in mice bearing established autologous NSCLC tumors. Furthermore, even after the loss of tumor-dominant TCR clonotypes and overgrowth of T cell clones that were marginally detectable in primary tumors during the *ex vivo* TIL expansion (25), we were capable of redirecting only TIL^STAb^ against EGFR tumor cells *in vitro* and *in vivo*.

It is worth noting the similarity between the dominant TCRβ CDR3 sequences of both post-REP products, and the finding that the overrepresented P1 ASSDGRYGYT clonotype is very similar to the ASDTGRYGYT clonotype that recognize the Epstein-Barr virus (EBV)-specific CD8 T cell epitope BRLF1_109-117_ (26), but also similar to the ASSEGSYGYT clonotype that recognizes the influenza A virus matrix protein 1 epitope M1_58–66_ (http://tools.iedb.org/tcrmatch/). Viral antigen-specific cells (EBV, cytomegalovirus, and influenza virus) have been described in TIL from lung and colorectal tumors (27), and a clonotype with the same TCRβ CDR3 sequence was enriched in lung tumor compared to normal lung tissue (28). One explanation for pathogen-specific TIL may be TCR cross-reactivity to both tumor and microbial antigens (27,29–31). Therefore, it is conceivable to assume that current *ex vivo* TIL generation protocols promote the expansion of virus-reactive TCR-bearing T cell clones present in primary lung tumors (32), with enhanced responsiveness/proliferative capacity and the dilution of NRTs.

Different approaches have demonstrated the efficacy of genetic modification during the *ex vivo* TIL expansion to enhance their antitumor capacity (8,16,33). Here, we demonstrate for the first time the feasibility of coupling lentiviral transduction and TIL expansion to generate a subpopulation of TCE-secreting T cells after a conventional REP. In a humanized PDX model, where EGFR^+^ NSCLC tumor fragments and TIL products from the same patient are sequentially transplanted into *hIL*-2 NOG transgenic mice, engineered TIL secreting an anti-EGFR TCE induce antitumor responses when administered intratumorally and after systemic administration. The autologous immune-humanized mouse model has been developed to predict TIL clinical responses in melanoma (19), but here we demonstrate its relevance for studying the therapeutic effects of ACT in epithelial tumors, such as lung cancer. NSCLC have a very heterogeneous tumor microenvironment (TME) (34) that acts as both a physical and chemical barrier, preventing T cell infiltration and reducing ACT potential. In fact, our study establishes a direct relationship between T cell infiltration of NSCLC tumor and antitumor response in the group treated with systemic TIL^STAb^ product. Although all NSCLC tumor fragments (5x5 mm) s.c. implanted in *hIL-2* NOG mice are derived from the autologous P1 tumor that was previously established and expanded in athymic nude mice, it is possible that the TME of the implanted fragments might be significantly different, compromising the ability of T cells to effectively penetrate the tumor. This is consistent with clinical studies in which therapeutic benefit depends on the ability of endogenous and/or adoptively transferred effector T cells to reach the target tumor tissue (35). Since tumor homing and infiltration by T cells is a key aspect to induce therapeutic responses after systemic administration of TIL^STAb^, it may be essential to improve the efficiency of lentiviral transduction to obtain TIL products with a higher percentage of TCE-secreting cells to induce a more efficient recruitment of intratumoral bystander T cells. Another option would be the generation of TIL^STAb^ expressing receptors for different chemokines secreted by tumor cells to improve TIL migration to tumor sites after re-infusion (8).

A major finding of this work is that tumor regression is mediated by CD4^+^ T cells. This is in consistent with recent observations of the cytotoxic potential of CD4^+^ cells (24,36–38), but the absence of class-II MHC molecules in cancer cells (**Supplementary Fig. S5A**) suggest that the CR found in TIL^STAb^-treated mouse could only have been driven by the local secretion of the anti-EGFR TCE. A significant number of scattered EGFP^+^ T cells could be detected in different areas of the tumor (**Fig. 3A** and **B**), most likely recruiting bystander cells, mainly CD4^+^ T cells with the dominant REP clonotype. When we analyzed the TCR repertoire of the tumor that experienced CR after systemic administration of TIL^STAb^ cells and compared to the TIL^NTd^-treated mouse with the highest CD3 infiltrate, we observed that the *ex vivo*-expanded ASSDGRYGYT clonotype was the predominant. Indeed, this ASSDGRYGYT clonotype was the only shared clonotype between P1 primary tumor and TIL^STAb^-treated mice. The predominant P1 TIL population is CD4^+^, hindering investigations of the effect of NRTs, which are present in the administered TIL product, on tumor growth after locally secreted TCE-mediated responses are triggered.

In conclusion, we have demonstrated the *in vitro* and *in vivo* antitumor activity of TCE- redirected CD4^+^ TIL^STAb^ bearing non-tumor dominant clonotypes, providing proof of concept for the possibility of genetically engineering TIL to secrete tumor-associated antigen-specific TCEs capable of redirecting T cells with non-tumor dominant clonotypes, regardless of their phenotype, for the treatment of solid tumors.

## AUTHORS’ DISCLOSURES

**Patient consent for publication.** Not applicable.

**Ethics approval and consent to participate.** All samples were obtained after written informed consent from the donors and all studies were performed according to the principles expressed in the Declaration of Helsinki and approved by the Institutional Research Ethics Committees of the hospitals and research centers involved. All the *in vivo* experiments were carried out in accordance with the CNIO Ethics Committee of Animal Experimentation. The experimental study protocols were additionally approved by local government (PROEX 084/15 and 253/19).

## AUTHORS’ CONTRIBUTIONS

Conceptualization, A.J.-R. and L.A-V; Methodology, A.J-R., A.G.-M., M.R-O., I.F., L.P-A., J.Z. and L.A-V; Formal analysis, A.J-R., M.M., C.C., E.M.G-M., A.N-B., R.T., E.P.; Investigation, A.J-R., M.M., C.C., B.B., E.M.G-M., D.N-A, C.D-A., A.R-F., L.D-A., A.N-B., J.L.R-P., I.F; Resources: A.G-M., M.R-O., L.P-A. and L.A-V; Writing - Original Draft: A.J-R., M.M., J.Z. and L.A-V.; Writing - Review & Editing, all authors; Funding acquisition, B.B., J.Z., L.P-A. and L.A-V.

## Supporting information

Inventory of Supporting Information

## ACKNOWLEDGMENTS

We thank Eduardo Cabello of the Radiophysics Service, and María Lasa-Lázaro, María José Castro-Panete and Esther Mancebo of the Immunology Service, all from the Hospital Universitario 12 de Octubre, for their technical assistance. We thank thr Histopathology Core Unit of the CNIO for technical support.

L.A-V. was supported by grants from the Spanish Ministry of Science and Innovation MCIN/AEI/10.13039/501100011033 (PID2020-117323RB-100, PDC2021-121711-100, CPP2022-009762 CPP2022-009765), the Instituto de Salud Carlos III (ISCIII) (DTS20/00089, PMPTA22/00167), the Comunidad de Madrid (P2022/BMD-7225 – NEXT- GEN-CART-MAD-CM), the Fundación CRIS contra el Cáncer (FCRIS-2021-0090, FCRIS- 2021-0090 and FCRIS-2024-0061), the Asociación Española contra el Cáncer (AECC) (PROYE19084ALVA, PRYGN234844ALVA), the Fundación ‘‘La Caixa’’ (HR21-0076-IL7R-LungCan) and the Fundación de Investigación Biomédica 12 de Octubre Programa Investiga (2022–0082). L.P-A. was supported by grants from the ISCIII (INGENIO, PMPTA22/00167, PI20/00870), the Comunidad de Madrid (P2022/BMD-7437–iLung) and the European Comission (TOPMESO Transcan), and the Fundación CRIS contra el Cáncer (2018/0043). J.Z. was supported by the ISCIII (PI20/01494) and the AECC (LABAE20044). B.B was supported by the ISCIII (PI20/01030). C.D-A. was supported by a predoctoral fellowship from the MCIN/AEI/10.13039/501100011033 (PRE2018-083445). L.D-A. was supported by a Rio Hortega fellowship from the ISCIII (CM20/00004).

## COMPETING INTERESTS

B.B., L.P-A and L.A-V are co-founders and shareholders of STAb Therapeutics, a spin-off company from the imas12. B.B. and L.A-V are inventors on the patent EP21708942 pending. L.D-A. and L.A-V are inventors on the patent EP23383410.0 pending. L.A-V is cofounder and shareholder of Leadartis, a company focused on unrelated interest. L.A-V reports speaker honoraria from MSD, Merck KGaA, BMS, Janssen, GSK, and Miltenyi, and receives grant support from Merck KGaA, all outside the submitted work. J.Z. has served as a consultant for Sanofi, Pfizer, Astra Zeneca, BMS, Novartis, NanoString, and Guardant Health; reports speaker honoraria from Pfizer, BMS, Roche, Astra Zeneca, NanoString and Guardant Health; and receives grant support from Astra Zeneca, Roche, and BMS, all outside the submitted work. L.P-A. is shareholder of Altum Squencing and STAb Therapeutics, has served as a consultant for Lilly, MSD, Roche, Pharmamar, Merck KGaA, Astra Zeneca, Novartis, Amgen, Pfizer, Sanofi, Bayer, BMS, Mirati, GSK, Janssen, Takeda, Regeneron, and Sanofi. L.P-A. received grant support from MSD, Astra Zeneca, BMS, Pfizer and Pharmamar, all outside the submitted work. R.T. and E.P. are consultants for the company NISOLAB.

