## Supplementary material for "CD4^+^ tumor-infiltrating lymphocytes secreting T cell-engagers induce regression of autologous patient-derived non-small cell lung cancer xenografts": Inventory of Supporting Information

Inventory of Supporting Information for

This Inventory includes:

- **List of Abbreviations**
- **Supplementary Figures**
- **Supplementary Tables**

#### LIST OF ABBREVIATIONS

**ACT**: Adoptive cell therapy

**CM**: central memory

**CR**: complete responses

**DCM**: DMEM complete medium

**DN**: double negative

**DP**: double positive

**DSP**: digital spatial profiling

**E:T**: Effector:target ratio

**EBV**: Epstein-Barr virus

**EGFP**: Enhanced green fluorescent protein

**EGFR LiTE**: anti-EGFR x anti-CD3 light T cell engager

**EGFR**: epidermal growth factor receptor

**ELISA**: enzyme-linked immunosorbent assays

**EM**: effector memory

**EMRA**: effector memory re-expressing CD45RA

**FC**: fold change

**FDR**: false discovery rate

**FFPE**: formalin-fixed and paraffin-embedded

**GZMB**: granzyme B

**H/E**: hematoxylin and eosin.

**hIL-2 NOG mice**: NOD.Cg-$Prkdc^{scid}$ $Il2rg^{tm1Sug}$ Tg(CMV-IL2)4-2Jic/JicTac

**IHC**: immunohistochemistry

**IL-2**: Interleukin-2

**KO**: knock out

**M**: mouse.

**NGS**: next-generation sequencing

**NRTs**: neoantigen reactive T cells

**NSCLC**: non-small cell lung cancer

**NTd**: Non-transduced

**P1**: Patient 1

**P2**: Patient 2

**PDX**: Patient-derived xenograft

**RCM**: RPMI complete medium

**REP**: rapid expansion protocol.

**rhIL-2**: recombinant human IL-2

**ROI**: regions of interest

**scFv**: single-chain variable fragment

**SD**: standard deviation

**SEM**: standard error of mean

**STAb**: Secreting T cell-engaging antibodies

**TAA**: tumor-associated antigen

**TCE**: T cell-engagers

**TCR**: T cell receptor

**TME**: tumor microenvironment

**TIL**: tumor-infiltrating lymphocytes

**Veh.**: vehicle

### SUPPLEMENTARY FIGURES

#### Supplementary Figure S1

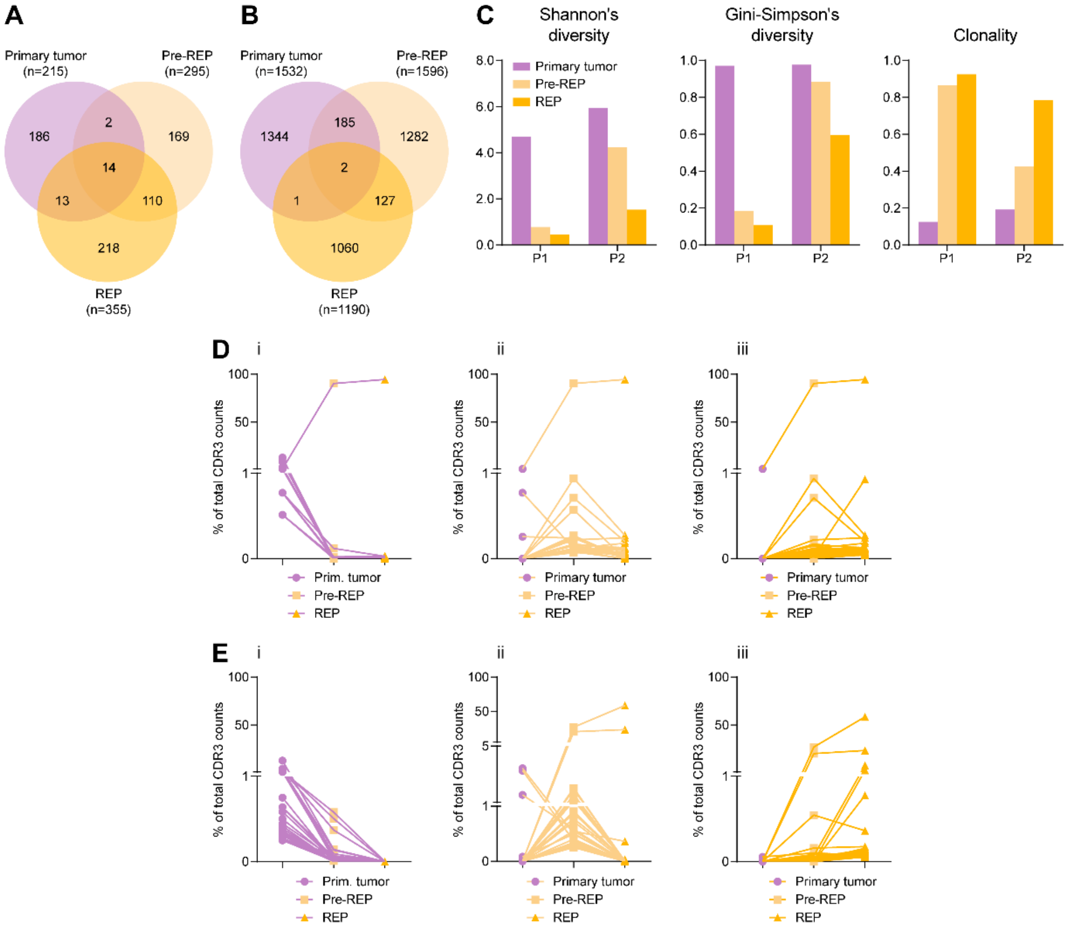


F

**P1 Primary tumor**

| TCRβ | # CDR3 aa | Counts |
|---|---|---|
| GPASVETRASISG | 13 | 50 |
| KEISSPLEII | 10 | 33 |
| LASGTAVKN | 9 | 11 |
| STLDFLDLQQH | 11 | 11 |
| EGTPYFRVQEGNEHWQNMY | 19 | 8 |
| RLLSLFEVLKTYFNFKGKA | 19 | 7 |
| ASSDGRYGYT | 10 | 4 |
| CHLPGSLADQRLMVRAR | 17 | 4 |
| GLRRYCRPQHRHHRQPLLS | 19 | 4 |
| IAGGEKRETWGQLPALWS | 18 | 4 |

**P1 Pre-REP**

| TCRβ | # CDR3 aa | Counts |
|---|---|---|
| ASSDGRYGYT | 10 | 10,908 |
| ASSQDPRAATYGYT | 14 | 113 |
| ASTDVAGGNQPQH | 13 | 86 |
| ASSEGAGYGYT | 11 | 69 |
| ASSEDRGSAGYT | 12 | 33 |
| ASSPGTGNYGYT | 12 | 31 |
| ASSQGVGAQH | 10 | 29 |
| TDPSIT | 6 | 29 |
| ASSEGRYGYT | 10 | 27 |
| ASSDGRYGYP | 10 | 20 |

**P1 REP**

| TCRβ | # CDR3 aa | Counts |
|---|---|---|
| ASSDGRYGYT | 10 | 46,078 |
| ASSIAGPLHEQF | 12 | 453 |
| ASSQDPRAATYGYT | 14 | 133 |
| ASSEGRYGYT | 10 | 119 |
| ASTDVAGGNQPQH | 13 | 98 |
| ARSDGRYGYT | 10 | 88 |
| AGSDGRYGYT | 10 | 63 |
| ASSGGRYGYT | 10 | 62 |
| ASSDGRYGYP | 10 | 57 |
| ASSDGRYGSP | 10 | 56 |

G

**P2 Primary tumor**

| TCRβ | # CDR3 aa | Counts |
|---|---|---|
| SNPHGNKRHVFH | 12 | 463 |
| LASGTAVKN | 9 | 163 |
| EVKEPV | 6 | 123 |
| GPASVETRASISG | 13 | 117 |
| STCLTI | 6 | 112 |
| KEISSPLEII | 10 | 63 |
| ASSLTSASGELF | 12 | 60 |
| PLPPLYKI | 8 | 55 |
| RLLSLFEVLKTYFNFKGKA | 19 | 40 |
| TESQNET | 7 | 27 |

**P2 Pre-REP**

| TCRβ | # CDR3 aa | Counts |
|---|---|---|
| ASSRDGQGQGEQF | 13 | 3,217 |
| ASSSGLYGYT | 10 | 2,411 |
| ASSRDRGGNQPQH | 13 | 240 |
| ASSLAGKNTEAF | 12 | 229 |
| ATSPGRGNQPQH | 12 | 185 |
| ATSLRDLNEKLF | 12 | 159 |
| ASSFKSGSGELF | 12 | 138 |
| ASSRTVNEKLF | 11 | 134 |
| ASSISGTGNEQF | 12 | 108 |
| ASSEGAAALH | 10 | 104 |

**P2 REP**

| TCRβ | # CDR3 aa | Counts |
|---|---|---|
| ASSRDGQGQGEQF | 13 | 122,322 |
| ASSSGLYGYT | 10 | 49,022 |
| ASSLQLVYQETQY | 13 | 16,389 |
| SVEPGGVTGELF | 12 | 5,966 |
| AISQGQGPYNEQF | 13 | 1,622 |
| ASSLGSLNTGELF | 13 | 759 |
| ASSRDGQGQGEQL | 13 | 367 |
| ASSRDGRGQGEQF | 13 | 327 |
| AGSRDGQGQGEQF | 13 | 306 |
| ASSRDGQGQGGQF | 13 | 274 |

**Figure S1. TCRβ clonotypes metrics and clonal dynamics during *ex vivo* TIL expansion. (A, B)** Shared clonotypes between primary tumors, pre- and post-REP TIL cultures from NSCLC P1 **(A)** and P2 **(B)**. **(C)** NSCLC P1 and P2 TCR Shannon's and Gini-Simpson's diversity scores, and TCR clonality scores in primary tumors and in pre-

and post-REP TIL cultures, calculated from total number of productive clonotypes with amino acid-length between 5-20. **(D, E)** Relative frequency of the TOP-30 largest clones of the primary tumor (i), the pre-REP (ii) or the post-REP (iii) CDR3 repertoire of samples from NSCLC P1 **(D)** and P2 **(E)**. (**F, G)** Detailed amino acid sequences of the TOP-10 largest clones of the primary tumor, the Pre-REP or the Post-REP CDR3 repertoire of samples from NSCLC P1 **(F)** and P2 **(G)**. P1, Patient 1; P2, Patient 2; REP, rapid expansion protocol.

#### Supplementary Figure S2

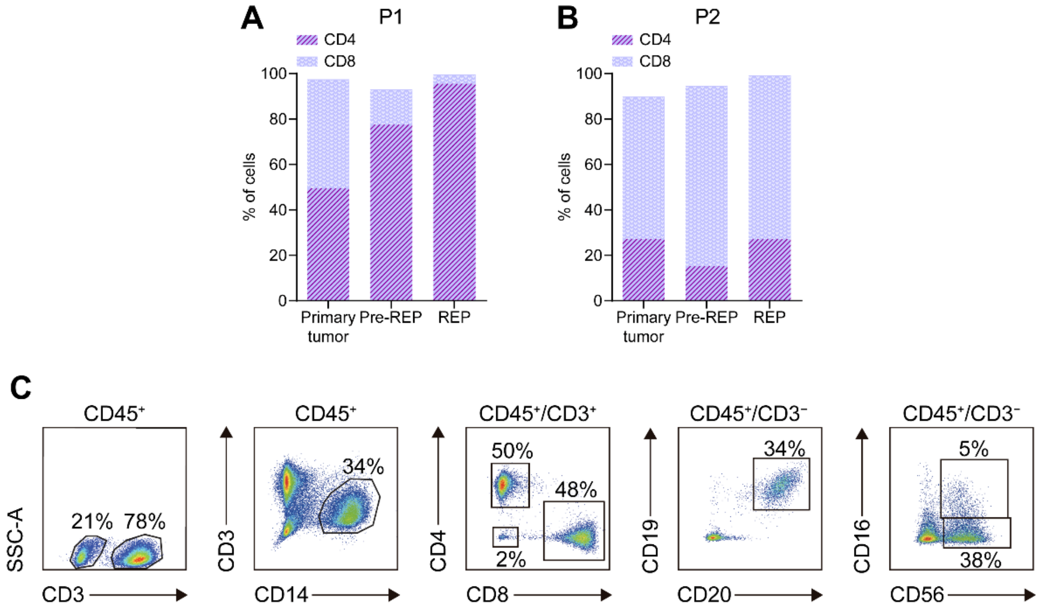


**Figure S2. Immune microenvironment phenotype in NSCLC patients. (A, B)** Percentages of CD4$^+$ and CD8$^+$ T cells from NSCLC P1 **(A)** and P2 **(B)**. **(C)** Percentages of total T (CD3$^+$), non-T (CD3$^–$), macrophages (CD3$^–$CD14$^+$), CD4$^+$ and CD8$^+$ T cells, B cells (CD19$^+$CD20$^+$) and NK cells (CD56$^+$CD16$^{+/–}$) within leukocytes (CD45$^+$) after P1 primary tumor disaggregation.

#### Supplementary Figure S3

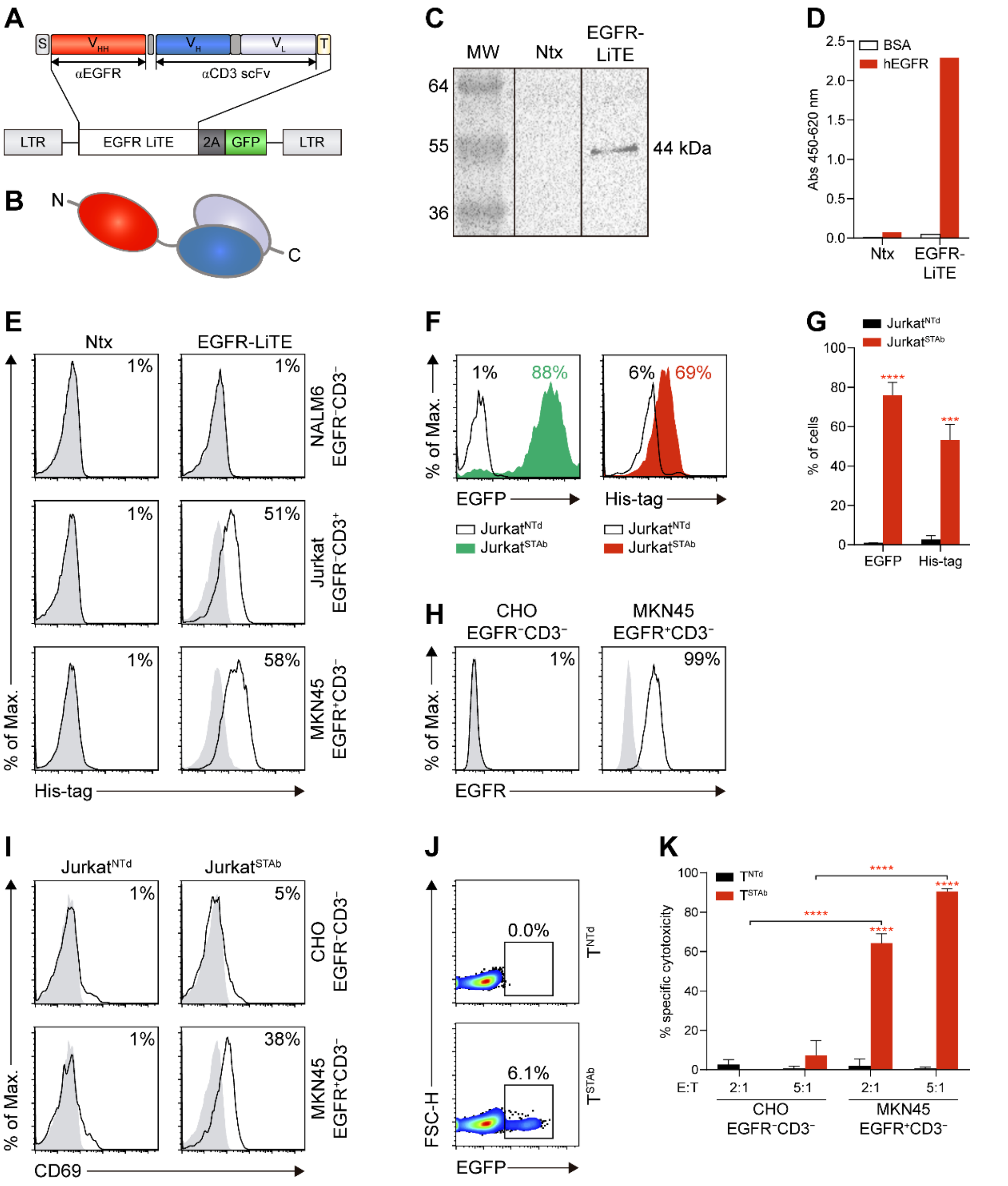


**Figure S3. Functionality of secreted anti-EGFR light T cell engagers (LiTE).** **(A, B)** Schematic diagrams showing the genetic **(A)** and domain structure **(B)** of the anti-EGFR x anti-CD3 Light T cell engager (EGFR LiTE) bearing the OncoM signal peptide (S, grey box), the anti-EGFR $V_{HH}$ gene (red box), the anti-CD3 scFv gene (blue boxes), and the Myc and His tags (light yellow box). The EGFR LiTE construct was cloned into a pCCL lentiviral-based backbone containing a T2A-EGFP cassette **(A)**. **(C, D)** Detection of soluble secreted EGFR LiTE in the conditioned media from pCCL transfected HEK293T cells by Western blot **(C)** or ELISA **(D)**. Conditioned media from non-transfected (NTx) cells was used as negative control. One representative experiment is shown. **(E)** Binding assays of conditioned media from NTx- or EGFR LiTE-transfected HEK293T cells to NALM6, Jurkat and MKN45 tumor cells. Specific binding was detected using anti-His-tag mAb and analyzed by flow cytometry. **(F)** Percentage of EGFP-positive (left) and

EGFR-LiTE decorated (right) engineered EGFR-STAb Jurkat ($Jurkat^{STAb}$) in comparison with non-transduced Jurkat cells ($Jurkat^{NTd}$). EGFR-LiTE decoration was detected using anti-His-tag mAb and analyzed by flow cytometry. **(G)** EGFP expression and EGFR-LiTE decoration analysis in $Jurkat^{NTd}$ and $Jurkat^{STAb}$ cells. Data represent mean+SEM of at least 3 independent experiments, and statistical significance was calculated by two-way ANOVA test corrected by Sidak's multiple comparisons test; $**p < 0.01$; $***p < 0.001$. **(H)** Cell surface expression profile of EGFR from target cells used in *in vitro* studies by flow cytometry. **(I)** T cell activation assay. $Jurkat^{NTd}$ or $Jurkat^{STAb}$ cells were co-cultured at a 1:1 E:T ratio with EGFR-negative (CHO) or EGFR-positive (MKN45) cells for 24 hours and CD69 expression analyzed by flow cytometry. **(J)** Percentage of EGFP-positive transduced primary T cells ($T^{STAb}$) in comparison with non-transduced primary T cells ($T^{NTd}$). **(K)** Specific cytotoxicity of $T^{NTd}$ or $T^{STAb}$ cells against EGFR-negative (CHO) or EGFR-positive (MKN45) cells at the indicated E:T ratios after 48 hours. The percentage of specific cytotoxicity was calculated by adding D-luciferin to detect bioluminescence. Data represent mean+SD and statistical significance was calculated by two-way ANOVA test corrected by Tukey's multiple comparisons test; $****p < 0.0001$. The inset numbers in **(E-J)** represent the percentage of cells staining positive for the indicated marker. EGFP, enhanced green fluorescence protein; SEM, standard error of the mean; SD, standard deviation.

#### Supplementary Figure S4

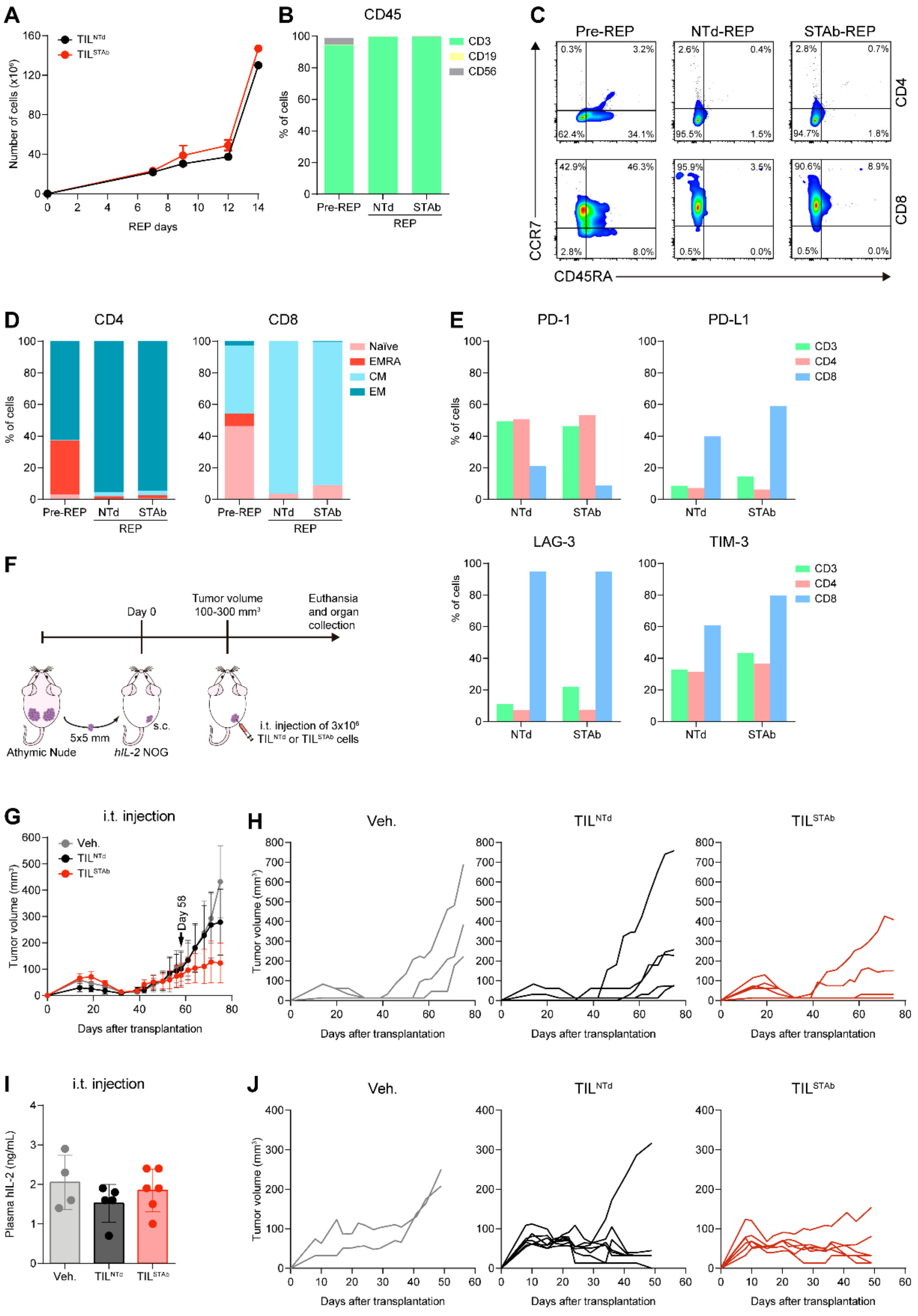

**Figure S4. Treatment of EGFR$^{+}$ NSCLC PDX mouse model with autologous engineered TCE-secreting TIL (cont.). (A)** Cell growth during the small-scale REP of non-transduced (TIL$^{NTd}$) and EGFR-STAb TIL (TIL$^{STAb}$). **(B)** Percentages of total T (CD45$^{+}$CD3$^{+}$), B (CD45$^{+}$CD19$^{+}$) and NK (CD45$^{+}$CD56$^{+}$) cells among Pre-REP TIL, TIL$^{NTd}$ REP and TIL$^{STAb}$ REP. **(C, D)** Plots **(C)** and graph barrs **(D)** representing the percentages of naïve, effector memory re-expressing CD45RA (EMRA), central memory

(CM), and effector (EM) $CD4^+$ or $CD8^+$ T cells among pre-REP, $TIL^{NTd}$ REP, and $TIL^{STAb}$ REP. **(E)** Exhaustion markers expression within total T, CD4 ($CD3^+CD4^+$) and CD8 ($CD3^+CD8^+$) cells after REP. **(F)** Experimental design of autologous PDX establishment. *hIL-2* NOG xenografted mice were intratumorally (i.t.) treated with $3\times10^6$ $TIL^{NTd}$ or $TIL^{STAb}$ cells when the average tumor volume reached approximately 100-300 $mm^3$. After tumors reached the indicated size, thirteen mice were randomized into three groups (3/5/5) and i.t. treated with Veh. (PBS supplemented with 300 IU/mL rhIL-2, $n$=3 mice), $TIL^{NTd}$ ($n$=5 mice) or $TIL^{STAb}$ ($n$=5 mice) cells. Black arrow indicates the day of the injection. Tumors sizes were measured with calipers. **(G, H)** Tumor growth curves represented as mean±SEM **(G)** or as individual mouse curves within the different groups **(H);** black arrow indicates the day of the i.t. injection. **(I)** Human IL-2 plasma levels in mice after TIL treatment (mean±SD). Significance was calculated by two-way **(G)** or one-way **(I)** ANOVA test corrected by Tukey's multiple comparisons test; no significant differences were found. **(J)** Tumor growth curves represented as individual mouse curves within the different groups of *hIL-2* NOG xenografted mice intravenously (i.v.) treated with $7.5\times10^6$ $TIL^{NTd}$ or $TIL^{STAb}$ cells when the average tumor volume reached approximately 100-300 $mm^3$. Veh., vehicle; SEM, standard error of mean; SD, standard deviation.

#### Supplementary Figure S5

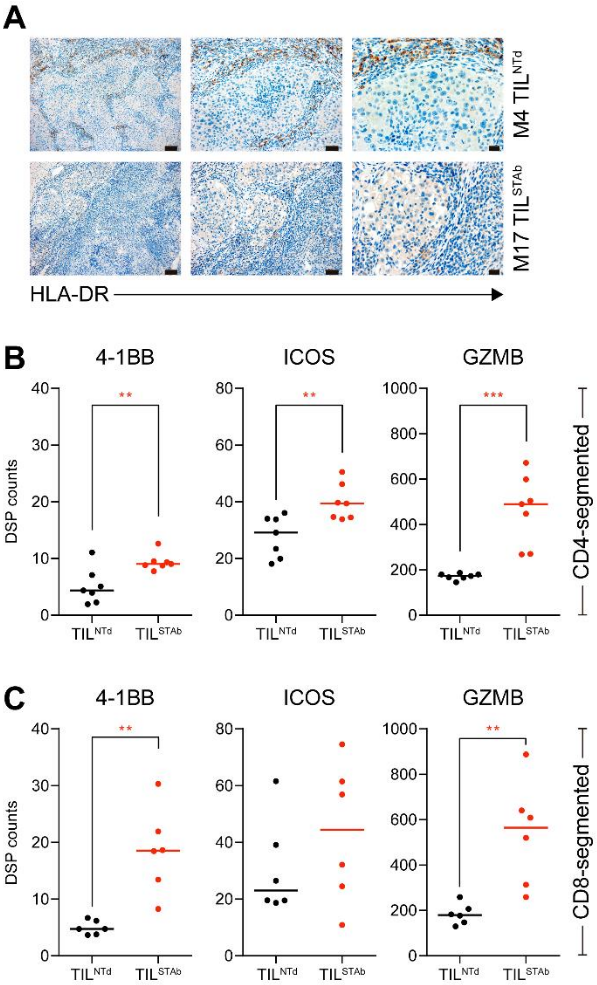


**Figure S5. Immunohistochemistry and DSP analysis of intravenously treated *hIL2* xenografted mouse 4 (TIL$^{NTd}$-treated) and mouse 17 (TIL$^{STAb}$-treated) (cont.). (A)** Expression of HLA-DR after treatment of *hIL-2* NOG xenografted mouse 4 (M4) with TIL$^{NTd}$ or mouse 17 (M17) with TIL$^{STAb}$. Scale bars: 100 µm (left), 50 µm (middle) and 20 µm (right). **(B)** Comparative analysis of 4-1BB, ICOS, and GZMB levels measured by DSP in in CD4 **(B)** or CD8 **(C)** compartments. Significance was calculated by an unpaired t-test. **p < 0.01; ***p < 0.001. GZMB, granzyme B.

#### Supplementary Figure S6

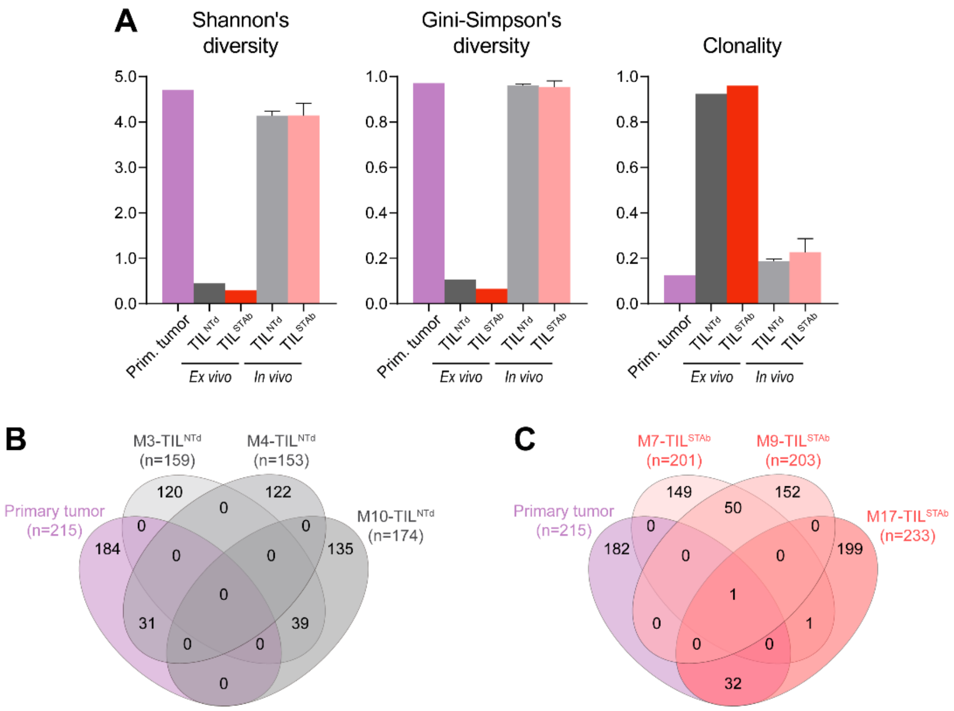


**Figure S6. TCR clonotypes composition of primary tumor and *in vivo* TIL-treated tumors from NSCLC P1. (A)** Shannon's and Gini-Simpson's diversity scores, and TCR clonality scores in primary tumor (Prim. Tumor), *ex vivo* expanded TIL and *in vivo* TIL-treated tumors. **(B, C)** Shared clonotypes between primary tumor and *in vivo* $TIL^{NTd}$-treated **(B)** or $TIL^{STAb}$-treated tumors **(C)**. Data in **(A-C)** are calculated from total number of unique productive clonotypes with 5-20 amino acid length CDR3 of TCRβ.

### SUPPLEMENTARY TABLES

#### Table S1. Antibodies

| Antigen | Clone | Host species / clonality | Brand | Reference | Application |
|---|---|---|---|---|---|
| Anti-GFP | D5.1 | Rabbit monoclonal | Cell Signaling Technology | 2956 | IHC |
| Anti-human CD3 | LN10 | Mouse monoclonal | Leica | CD3-565-L-CE | IHC |
| Anti-human CD3 FLEX-X | – | Rabbit Polyclonal | Dako | IR503 | IHC |
| Anti-human CD4 | 4B12 | Mouse monoclonal | Leica | CD4-368-L-CE | IHC |
| Anti-human CD8 | 4B11 | Mouse monoclonal | Leica | CD8-4B11-L-CE | IHC |
| Anti-human CD31 | JC70A | Mouse monoclonal | Leica | CD31-607-L-CE | IHC |
| Anti-human CD31 | EPR17259 | Rabbit monoclonal | Abcam | ab182981 | IHC |
| Anti-human EGFR | H11 | Mouse monoclonal | Dako | M3563 | IHC |
| Anti-human HLA-DR, α-chain | TAL.1B5 | Mouse monoclonal | Dako | M0746 | IHC |
| Anti-human Perforin | 5B10 | Mouse monoclonal | Invitrogen | MA512469 | IHC |
| Anti-human FOXP3 | 236A/E7 | Mouse monoclonal | eBioscience | 14-4777-82 | IHC |
| Anti-His APC | GG11-8F3.5.1 | Mouse monoclonal | Miltenyi Biotec | 130-119-782 | FC |
| Anti-human CD3 V450 | UCHT1 | Mouse monoclonal | BD Horizon | 560365 | FC |
| Anti-human CD3 FITC | HIT3a | Mouse monoclonal | BD Pharmingen | 561802 | FC |
| Anti-human CD4 BUV395 | RPA-T4 | Mouse monoclonal | BD Horizon | 564724 | FC |
| Anti-human CD4 PerCP-Cy5.5 | SK3 | Mouse monoclonal | BD | 332772 | FC |
| Anti-human CD8 BUV563 | RPA-T8 | Mouse monoclonal | BD Horizon | 565695 | FC |
| Anti-human CD8 APC-Cy7 | SK1 | Mouse monoclonal | BioLegend | 344714 | FC |
| Anti-human CD14 BV605 | M5E2 | Mouse monoclonal | BD Horizon | 564054 | FC |

| | | | | | |
|---|---|---|---|---|---|
| Anti-human CD16 Brilliant Violet 785™ | 3G8 | Mouse monoclonal | Biolegend | 302045 | FC |
| Anti-human CD19 PE-Cy7 | HIB19 | Mouse monoclonal | eBioscience™ | 25-0199-42 | FC |
| Anti-human CD19 APC | HIB19 | Mouse monoclonal | BD Pharmingen | 555415 | FC |
| Anti-human CD20 APC | 2H7 | Mouse monoclonal | BioLegend | 302309 | FC |
| Anti-human CD25 FITC | M-A251 | Mouse monoclonal | BD Pharmingen | 555431 | FC |
| Anti-human CD45 V500-C | 2D1 | Mouse monoclonal | BD Horizon | 655873 | FC |
| Anti-human CD45 Alexa Fluor® 700 | HI30 | Mouse monoclonal | BD Pharmingen | 560566 | FC |
| Anti-human CD45RA V500 | HI100 | Mouse monoclonal | BD Horizon | 561640 | FC |
| Anti-human CD56 PE-Cy7 | NCAM16.2 | Mouse monoclonal | BD | 335826 | FC |
| Anti-human CD56 APC | B159 | Mouse monoclonal | BD Horizon | 555518 | FC |
| Anti-human CCR7 BV421 | 150503 | Mouse monoclonal | BD Horizon | 562555 | FC |
| Anti-human EGFR | EGFR.1 | Mouse monoclonal | BD Horizon | 566254 | FC |
| Anti-human HLA-DR PE-Cy7 | G46-6 | Mouse monoclonal | BD Pharmingen | 560651 | FC |
| Anti-human TCRg/d FITC | 11F2 | Mouse monoclonal | BD | 347903 | FC |
| Anti-human TCRa/b PE | BW242/412 | Mouse monoclonal | Miltenyi Biotec | 130-113-531 | FC |
| Anti-human CD4 Alexa Fluor® 594 | EPR6855 | Rabbit monoclonal | Abcam | ab277931 | DSP |
| Anti-human CD8 Alexa Fluor® 647 | C8/144B | Mouse monoclonal | Novus Biologicals | NBP2-34588AF647 | DSP |
| Anti-His | PentaHis | Mouse monoclonal | QIAGEN | 34660 | ELISA |
| Anti-mouse IgG (H+L) HRP | – | Goat polyclonal | Jackson ImmunoResearch | 115-035-166 | ELISA |

| | | | | | |
|---|---|---|---|---|---|
| Anti-Myc | 9E10 | Mouse monoclonal | Merck Millipore | 05-419 | WB |
| Anti-mouse IgG (Fc specific) HRP | – | Goat polyclonal | Sigma-Aldrich | A2554 | WB |

IHC, Immunohistochemistry; FC, Flow Cytometry; DSP, Digital Spatial Profiling; ELISA, Enzyme-linked immunosorbent assays; WB, Western blot.

#### Table S2. Rest of reagents

| Antigen | Brand | Reference | Application |
|---|---|---|---|
| Neutral buffered formalin | Sigma-Aldrich | HT501128 | IHC |
| DNP-labeled Alu positive control probe II | Ventana, Roche | 05272041001 | ISH |
| DAPI | Sigma-Aldrich | D9542-10MG | FC |
| FcR Blocking Reagent, human | Miltenyi Biotec | 130-059-901 | FC |
| LIVE-DEAD Fixable Blue Dead Cell Stain Kit | eBioscience | L34961 | FC |
| LIVE-DEAD Fixable Violet Dead Cell Stain Kit | eBioscience | L34963 | FC |
| Anti-human CD45 - GeoMx Solid Tumor TME Morphology marker | NanoString Technologies | GMX-PRO-MORPH-HST-12 | DSP |
| Anti-human PanCk - GeoMx Solid Tumor TME Morphology marker | NanoString Technologies | GMX-PRO-MORPH-HST-12 | DSP |
| SYTO 13 Nucleic Acid Stain - GeoMx Solid Tumor TME Morphology marker | NanoString Technologies | GMX-PRO-MORPH-HST-12 | DSP |
| GeoMx protein Slide prep kit PCLN | NanoString Technologies | 121300312 | DSP |
| GeoMx Immune Cell Profiling Panel | NanoString Technologies | GMX-PROCO-NCT-HICP-12 | DSP |
| GeoMx IO Drug Target Panel | NanoString Technologies | GMX-PROMOD-NCT-HIODT-12 | DSP |
| GeoMx Immune Activation Status Panel | NanoString Technologies | GMX-PROMOD-NCT-HIAS-12 | DSP |
| GeoMx Hyb Code Pack Protein | NanoString Technologies | GMX-PRO-HYB-96 | DSP |
| nCounter Master Kit | NanoString Technologies | NAA-AKIT-012 | DSP |
| Image-iT™ Fixative Solutions | Invitrogen™ | FB002 | DSP |
| Citrate Buffer, pH 6.0, 10x, Antigen Retriever | Sigma-Aldrich | C9999 | DSP |
| TintoRetriever Pressure Cooker | Bio SB | 7008 | DSP |
| EGFR Fc chimera protein | R&D Systems | 344-ER | ELISA |
| Nunc MaxiSorp™ | ThermoFisher Scientific | M9410-1CS | ELISA |
| Tween®20 | Sigma-Aldrich | P1379 | ELISA |
| Tetramethylbenzidine | Sigma-Aldrich | T0440 | ELISA |
| Novex™ WedgeWell™ 10–20% Tris-glycine gels | Invitrogen™ | XP10202BOX | WB |
| Immobilon-PVDF Transfer Membrane | Merck Millipore | IPVH00010 | WB |
| Pierce™ ECL Plus Western Blotting Substrate | ThermoFisher Scientific | 32132 | WB |

| | | | |
|---|---|---|---|
| QIAamp DNA Blood Mini Kit | QIAGEN | 50951104 | NGS |
| FFPE RNA/DNA Purification Plus Kit | Norgen Biotek | 54300 | NGS |
| SuperPlex™ Premix | Takara | 638543 | NGS |

IHC, Immunohistochemistry; ISH, *In situ* hybridization; FC, Flow Cytometry; DSP, Digital Spatial Profiling; ELISA, Enzyme-linked immunosorbent assays; WB, Western blot.

#### Table S3. TCRβ statistics

| Patient | Sample | Total queries | Total unique clonotypes | Total reads assigned to unique clonotypes | Total unique productive clonotypes | Total reads assigned to unique productive clonotypes | Total unique productive clonotypes with 5-20 aa CDR3 | Total reads assigned to unique productive clonotypes with 5-20 aa CDR3 | Shannon Index, H | Gini-Simpson Index, D | Pielou's Evenness, J | Clonality score, C |
|---|---|---|---|---|---|---|---|---|---|---|---|---|
| P1 | Primary tumor | 186,710 | 3,586 | 6,684 | 487 | 753 | 215 | 389 | 4.702 | 0.971 | 0.876 | 0.124 |
| | Pre-REP | 57,164 | 2,084 | 20,200 | 488 | 12,204 | 295 | 12,085 | 0.772 | 0.185 | 0.136 | 0.864 |
| | REP | 131,062 | 3,171 | 68,505 | 720 | 49,020 | 355 | 48,733 | 0.447 | 0.106 | 0.076 | 0.924 |
| P2 | Primary tumor | 639,696 | 13,166 | 42,613 | 2,760 | 6,809 | 1,532 | 3,597 | 5.931 | 0.977 | 0.809 | 0.191 |
| | Pre-REP | 320,404 | 10,625 | 53,727 | 2,839 | 13,776 | 1,596 | 11,896 | 4.237 | 0.884 | 0.574 | 0.426 |
| | REP | 462,332 | 3,096 | 286,422 | 1,913 | 209,019 | 1,190 | 209,005 | 1.529 | 0.595 | 0.216 | 0.784 |

P1, patient 1; P2, patient 2; REP, rapid expansion protocol; aa, amino acid. Shannon's and Gini-Simpson's indexes, Pielou's Evenness and Clonality scores were calculated from total number of productive clonotypes with CDR3 size between 5-20 aa.
